# Brain connectivity-informed regularization methods for regression

**DOI:** 10.1101/117945

**Authors:** Marta Karas, Damian Brzyski, Mario Dzemidzic, Joaquin Goni, David A. Kareken, Timothy W. Randolph, Jaroslaw Harezlak

## Abstract

A challenging problem arising in brain imaging research is principled incorporation of information from different imaging modalities. Frequently each modality is analyzed separately using, for instance, dimensionality reduction techniques which result in a loss of mutual information. We propose a novel regularization method to estimate the association between the brain structure features and a scalar outcome within the linear regression framework. Our regularization technique provides a principled approach to utilizing external information arising from the structural brain connectivity to inform the estimation of the regression coefficients. Our proposal extends the classical Tikhonov regularization framework by defining a penalty term based on the structural connectivity-derived Laplacian matrix. In the work presented, we address both theoretical and computational issues. The approach is illustrated using simulated data and compared with other penalized regression methods. Finally, we apply our regularization method to study the associations between the alcoholism phenotypes and brain cortical thickness using a diffusion tensor imaging (DTI) derived measure of structural connectivity.

## 1 Introduction

In vivo brain imaging studies usually collect multiple imaging data types, but most often the analysis is done for each data type separately, which does not adequately take into account the brain's complexity. Statistical methods that simultaneously utilize and combine multiple data types can provide a more holistic view of brain (dys)function. We propose a novel statistical methodology that combines imaging data to derive a more complete picture of disease markers. In particular, we rigorously model associations between scalar phenotypes and imaging data while incorporating prior scientific knowledge. Specifically, we incorporate structural connectivity measures to model the association between the brain cortical thickness and alcoholism-related phenotypes. However, our methodology is more general and applicable to a variety of continuous outcomes and connectivity measures.

We work with a linear regression model where the scalar outcome for each subject *i, y_i_* (vector of outcomes **y**) is modeled as a linear combination of covariates *x*_1_,… *,x_m_* (matrix of covariates, *X*) whose contribution is not penalized and predictors *z*_1_*,…,z_p_* (matrix of predictors, *Z*) whose contribution is penalized. Optimization problem for the parameter estimation can be written as

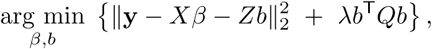

where the estimates of *β* and *b* are the best linear unbiased estimators (BLUE) and the best linear unbiased predictors (BLUP), respectively; *Q* incorporates the penalization information and λ is regularization parameter.

The general idea of incorporating structural information into regularized methods is well established (see e.g. Bertero and Boccacci (1998), Engl et al. (2000), Phillips (1962)). Such information is provided by a matrix that is inserted into the penalty term and constructed depending on the application. One of the most commonly used matrices are the second–difference matrices, which impose smoothness on the estimates (Huang et al., 2008). Hastie et al. (1995) note that under the situation with many highly correlated predictors it is “efficient and sometimes essential to impose a spatial smoothness constraint on the coefficients, both for improved prediction performance and interpretability.” A more general way of viewing the problem is that the modified penalty should take into account some presumed structure (a priori association) in the signal (Tibshirani and Taylor, 2011; Slawski et al., 2010). Such presumed structure can be represented mathematically in terms of an adjacency matrix which shows strength of connections between variables (corresponding to nodes of the graph), and is reflected in the penalty term via Laplacian (Chung, 2005). This procedure constitutes the basis for approaches presented by Li and Li (2008) and Randolph et al. (2012).

Prior information represented by a matrix *Q* may be incorporated into a regression framework using various types of penalty terms. Tikhonov regression (Tikhonov, 1963) and PEER (Partially Empirical Eigenvectors for Regression) (Randolph et al., 2012), employ the penalty of the form λ*b^T^Qb,* with a symmetric semipositive definite matrix *Q*. The choice *Q* = *I* results in an ordinary ridge regression, which is the most commonly used method of regularization of the ill-posed problems. The connection of this type of penalty with *l*_1_ norm was analyzed by Li and Li (2008) and Slawski et al. (2010). The penalized version of a linear discriminant analysis (LDA) was considered by Hastie et al. (1995), while Tibshirani and Taylor (2011) proposed the *l*_1_ norm imposed on the matrix Q times the coefficient vector. In each of these, different choices of *Q* give rise to a variety of well–known models, including the fused lasso (Tibshirani et al., 2005).

The natural problem that arises with penalized methods is the selection of a regularization parameter. Standard techniques to address this issue include the L–curve criterion and either cross-validation (CV) or generalized cross-validation (GCV) (see Craven and Wahba (1979), Hansen (1998), Brezinski et al. (2003)). In Randolph et al. (2012), authors take an advantage of the equivalence between the considered optimization problem and the Restricted Maximum Likelihood (REML) estimation (Ruppert et al., 2003; Maldonado, 2009) in the linear mixed model (LMM) framework. GCV and REML proceed by optimizing function of λ. According to experiments performed by P. T. Reiss (2009), GCV more likely finds a local, but not global, optimum of this function as compared to REML. Better performance of REML was also confirmed in simulations performed in this article and, consequently, the equivalence with LMM was chosen as a technique for selection of regularization parameters in our proposal.

The linear mixed model equivalence with the penalized problem of Tikhonov type, with penalty λ*b^T^Qb,* leads to the assumption that the prior distribution of *b* is of the form 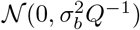, for some σ^2^_*b*_ > 0. This also shows a connection with the Bayes approach (see e.g. Maldonado (2009)) and raises the interpretation problem in the situation when *Q* is a singular matrix, which may be the case in some applications. This concern needs to be addressed, since the reduction to the ridge regression, which is the first step in λ selection procedure via REML, requires the invertibility of *Q*. One possible solution is to use the *Z*-weighted generalized inverse of *Q*, (Elden, 1982; Hansen, 1998), which is defined based on both the penalty matrix *Q* and the design matrix *Z*. This solution produces the assumed distribution for *b* which is not “purely prior”, since it depends on a specific dataset. Methods introduced in this work provide an alternative approach of handling the non-invertibility of the matrix *Q*. In our approach, we take an alternative point of view for the prior distribution of *b*; we assume that the unknown true or optimal variance–covariance, ℚ, of *b* is (potentially) non-singular and that a singular matrix *Q*, defined by the user, is “close” to the ℚ. This reasoning prompts us to accept that there exists an entire set of matrices which carry some amount of information about the true correlation structure of *b* (i.e. a set of informative matrices) and by using any of them we should be able to obtain a more accurate estimation and prediction than in the case when the knowledge about the signal structure is not available. In the next step, we assume that by adding the identity matrix multiplied by relatively small constant to matrix *Q* (“riPEER-c” approach) or by not penalizing the eigenvectors associated with the zero eigenvalues of the penalty matrix (“vrPEER” approach), the amount of information it carries will not change significantly. Finally, since it is not obvious how large the multiplicative constant should be, our final proposed method, riPEER, uses REML to estimate both penalty parameters simultaneously.

riPEER can be viewed as a special case of the extension of the general-form Tikhonov regularization, known as the multi-parameter Tikhonov problem (Belge et al., 2002; Brezinski et al., 2003; Lu and Pereverzev, 2011). Multi-parameter Tikhonov problem considers *k* matrices, *Q*_1_,…,*Q_k_*, together with *k* parameters λ_1_,…,λ_*k*_, and defines the penalty as 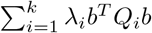. The set of *k* regularization parameters must be selected and a few ideas, such as a higher dimension L–curve (Belge et al., 2002), the discrepancy principle (Lu and Pereverzev, 2011), and an extension of GCV (Brezinski et al., 2003) were previously proposed. To the best of our knowledge, our work is the first to use the equivalence of the multi–parameter Tikhonov problem with the linear mixed model framework to estimate the regularization parameters.

To define the measure of similarity between the brain regions, we used the density of connections calculated from the measurements obtained from the DTI scans. The similarity measure between the regions *i* and *j* constitutes the (*i,j*)-entry of the symmetric adjacency matrix 𝒜, which is used to construct the normalized Laplacian, described in detail in Section 2.2. We exploit this structure as prior scientific knowledge in our proposed regularization methodology. A sample graph, its corresponding adjacency matrix, and a heatmap of the normalized Laplacian are presented in Figure 1.

**Figure 1:**
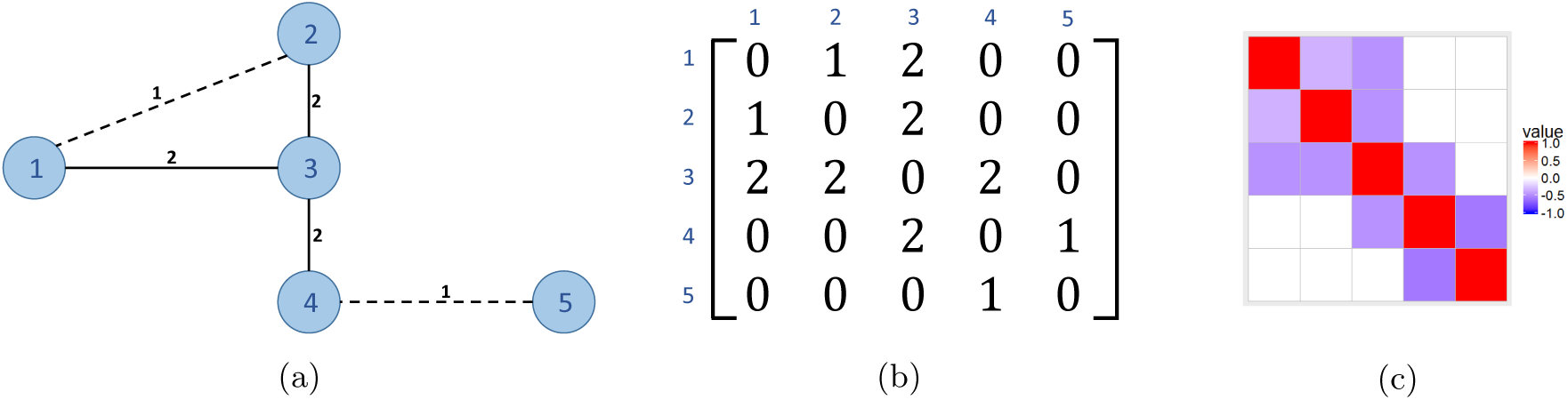
(a) Connectivity graph with five nodes and node similarity expressed as edge weights; (b) the corresponding adjacency matrix 𝒜 and (c) the heatmap of the normalized Laplacian.

The remainder of our article is organized as follows. In Section 2, we formulate our statistical model, propose the estimation procedure and discuss their main properties. We also describe the penalty term construction employing graph theory concepts and regularization parameter selection. Our new methods are described in Section 3 and simulations to study their validity are reported in Section 4. Finally, in Section 5, we apply our methodology to study the association of the brain’s cortical thickness and alcoholism phenotypes. The conclusions and a discussion are summarized in Section 6.

## 2 Statistical Model and Penalization

Motivated by the brain imaging data applications, we consider a situation where the covariates and penalized predictors are distinct. For example, information about the connectivity between the cortical brain regions is used to penalize only the predictors corresponding to cortical measurements, whereas other variables, such as demographic covariates and overall intercept, are included in the regression model as unpenalized terms.

### 2.1 Statistical model

Consider the general situation, where we have *n* observations of a random variable, stored in vector *y*, and the design matrix, [*Z X*], is given, where *Z* and *X* are n × *p* and *n* × *m* design matrices, respectively. Here, *p* denotes the number of all penalized variables and *m* is a number of unpenalized covariates. We assume that the unknown vectors *b* and *β* satisfy the multiple linear regression model,

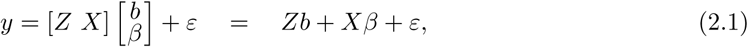

where *ε* ~ 𝒩(0, σ^2^*I_n_*) is the vector of random errors.

Here, *X* is the *n* by *m* design matrix containing unpenalized covariates or, equivalently, columns corresponding to variables for which the connectivity information is not given. The standard procedure is to mean-center all columns in the design matrix to remove the intercept. In such case, *X* can be an empty matrix (i.e. the situation with *m* = 0 is taken into consideration) and then model (2.1) takes the form *y* = *Zb* + *ε*. The matrix *Z* corresponds to the predictors for which the connectivity information is available. Such information is given by a *p* by *p* symmetric matrix *A* (also known as the connectivity matrix) which can be treated as a prior knowledge and employed in the estimation, e.g. via a Laplacian matrix *Q* (see Section 2.2), which can then be incorporated into the penalty term.

### 2.2 Defining *Q* as a normalized Laplacian

In this work, we use the normalized Laplacian matrix, see Chung (2005). Let *A* = [*a_ij_*], 1 ≤ *i,j* ≤ *p* be an adjacency matrix (symmetric matrix with non-negative entries and zeros on the diagonal), which defines the strength of the connections between nodes. The diagonal entries of the (unnormalized) Laplacian are *d_i_* := ∑_*j≠i*_, *a_ij_*, representing the sum of all weighted edges connected to node *i*.

The normalized Laplacian is defined as the *p* by *p* matrix *Q*:

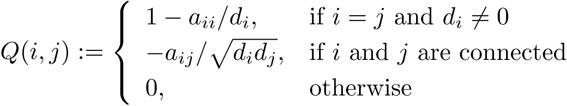

It is worth noting that a Laplacian is a symmetric matrix. The justification why a matrix constructed in such a way could be incorporated in the penalty term is explained by the following property (see e.g. Li and Li (2008)). If we let *Q* to be the normalized Laplacian then

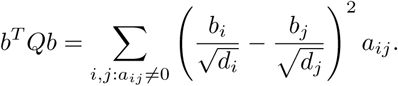

The result above shows that defining penalty as λ*b^T^ Qb* implies that vectors that differ too much over linked nodes get more penalized. Scaling by 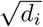 allows a small number of nodes with large *d_i_* to have more extreme values. The proposition above also implies that for any adjacency matrix *A*, Laplacian is a positive semi-definite matrix. To determine that *Q* has zero as the smallest eigenvalue, it is enough to consider a *p* dimensional vector, *b̃*, defined as 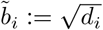 Then *b̃*^*T*^*Qb̃* = 0.

### 2.3 Estimate and its properties

If *Q* is a symmetric and positive definite matrix (hence invertible), we can use the estimate obtained as the solution to a convex optimization problem of the general-form Tikhonov regularization, i.e.,

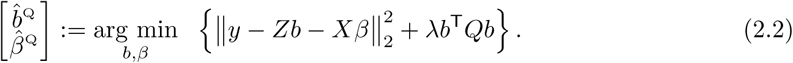

Here λ > 0 is a tuning parameter which can be estimated using the equivalence with a corresponding linear mixed model. This estimation procedure is implemented in Randolph et al. (2012). In our case, there is clear distinction between the penalized – Z and unpenalized – *X* covariates. By defining 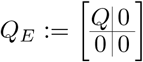, the minimization procedure (2.2) is equivalent to

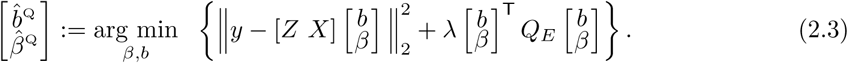

The solution to the above problem can be analytically found and is given by

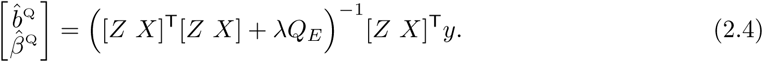

Randolph et al. (2012) describe how the generalized singular value decomposition (GSVD) (see also Hansen (1998), Golub and Van Loan (2013), Bjorck (1996), Paige and Saunders (2006)) is a useful tool for understanding the role played by the *l*^2^ penalization terms in a regularized regression model. In particular, GSVD provides a tractable and interpretable series expansion of the estimate in (2.4) in terms of the generalized eigenvalues and eigenvectors.

### 2.4 Regularization parameter selection

We consider two parameter selection procedures, cross-validation and REML. Simulations performed in Section 4 show the advantage of the later over the former. Therefore, in our final procedure, the equivalence with linear mixed model is used to obtain λ.

We briefly review here the basics of the LMM theory necessary to present our proposal (see e.g. Demidenko (2004) and C. E. McCulloch (2008)). Here, we consider the multiple linear regression model, *y* = *Xβ* + *Zb* + *ε*, with uncorrelated random vector *ε*, i.e. we assume that 𝔼(*ε_i_*, *ε_j_*) = 0 for *i* ≠ *j*. To present the idea we begin with the case where *Q* has an inverse that matches (up to a multiplicative constant) the variance-covariance matrix of random effects. In summary, we assume the following:

A.1 *β* is the vector of fixed effects and *b* is vector of random effects,
A.2 𝔼(*b*) = 0 and 𝔼(*ε*) = 0,
A.3 *cov*(*b*) = σ^2^*_b_Q*^−1^ and *Cου*(*ε*) = σ^2^*Ι_n_*, for some unknown σ^2^*_b_* > 0 and σ^2^ > 0,
A.4 *b* and *ε* are uncorrelated.

As noted above, knowing *Q* corresponds to knowing the correlation matrix of *b* up to the multiplicative constant, σ^2^_*b*_ > 0, which can be interpreted as a measure of signal amplitude. By assuming two-, three- and higher parameter families, one can generalize this to the case where knowledge about the correlation is less rich. In the extreme case no additional conditions for *Cov*(*b*) are assumed, other than that it is a symmetric, positive definite matrix, which results in (*p*^2^ + *p*)/2- parameter family of matrices.

Under the model (2.1) with assumptions A.1-A.4, it can be shown that 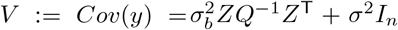. The best linear unbiased estimator (BLUE) of *β* and best linear unbiased predictor (BLUP) of *b* are given by the following equations

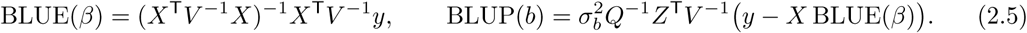

Equivalently the expressions in (2.5) can also be obtained as the solution to an optimization problem. Assume that *b* has multivariate normal distribution: *y*|*b* ~ 𝒩 (*Xβ* + *Zb,σ*^2^*I_n_*) and 𝒩(0,σ^2^_*b*_ *Q*^−1^).If σ^2^_*b*_ σ^2^ are known, this yields the following log-likelihood function

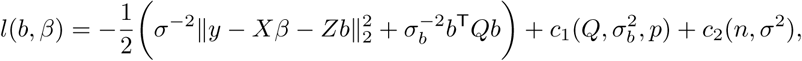

where functions *c*_1_ and *c*_2_ do not depend on either *b* or *β*. Looking for the maximum likelihood estimates simply leads to the problem (2.2), with λ:= σ^2^/σ^2^_*b*_. It can be shown that the optimal values of *β* and *b* are exactly given by BLUE and BLUP defined in (2.5). Our proposal provides an objective and statistically rigorous way to choose the tuning parameter λ in (2.2) when *Q* is a symmetric and positive definite matrix. Indeed, we can use REML to obtain the estimates of σ^2^ and σ^2^/σ^2^_*b*_ in the model characterized by A.1 – A.4 and define λ̂ as σ̂^2^/σ̂^2^_*b*_.

The first step is the *conversion* of the optimization problem to the ordinary ridge regression. Since *Q* is a symmetric, positive-definite matrix it can be decomposed as *Q* = *L*^T^*L*, where *L* is an invertible matrix. Now

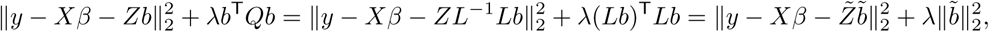

for *Z̃* := *ZL*^−1^ and *b̃* := *Lb,* which allows us to assume that *Q* = I in A.3 without loss of generality. This conversion procedure changes the general Tikhonov formulation to the ordinary ridge regression. One of the clear advantages of the the ordinary ridge is that it is easily implemented in a variety of existing software packages that support the LMM framework.

### 2.5 Statistical inference

We utilize a testing framework proposed in Ruppert et al. (2003) to test for the significance of the penalized regression coefficients *b* and provide the (1 – *α*) confidence intervals for the riPEER estimate via

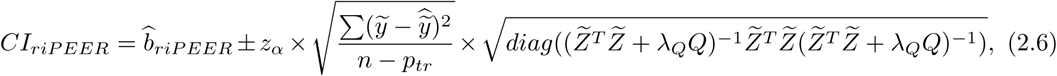

where *z_α_* is the value of the inverse of CDF of the standard normal distribution in 1 – *α*/2 and *p_tr_* is the trace of a “hat” matrix *H* given by:

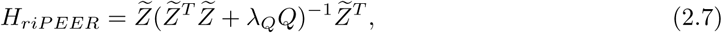

where *ỹ*, *Z̃* denote a vector of response variable *y* and penalized predictors data matrix *Z*, respectively (after regressing out the nonpenalized variables).

## 3 Regularization Methods

The standardization procedure described in Section 2 can be applied when *Q* is a symmetric positive definite matrix. However, the normalized Laplacian is always a singular matrix. Thus, the methods described cannot be readily applied.

Suppose that *Q* in (2.2) is non-invertible and has rank equal to *r* with *r < p.* Since matrix Q is singular, we can not simply assume that *b* ~ 𝒩(0,σ^2^_*b*_*Q*^−1^), as in A.3. However, it is reasonable to assume the existence of an unknown matrix ℚ such that *b* ~ 𝒩(0,σ^2^_*b*_ℚ^−1^) which is not identical, but, in some sense, close to *Q*. The intuition is that by making a “small modification” to *Q*, targeted to remove the singularity, one may produce an invertible matrix which is still “close” to *Q* and can use the information that it carries as described Section 2. In this section, we propose three approaches to address the singularity of the regularization matrix *Q*.

### 3.1 Constant riPEER (riPEER-c)

The simple and natural idea of modifying *Q* is by defining *Q̃* := *Q* + λ_2_*I_p_* for some fixed parameter λ_2_. We refer to this method as Constant riPEER (riPEER-c), where specific steps are presented below

#### Algorithm 1 riPEER-c

**input:** matrices *Z, X*, positive semi-definite matrix *Q,* λ_*2*_ > 0

**1:** define *Q̃* := *Q* + λ_2_*I_p_*;

**2:** decompose *Q* as *Q* = *L*^T^*L*;

**3:** find REML estimates *σ̂* and σ̂^2^, of σ^2^ and σ̂^2^*_b_*, σ^2^ and σ̂^2^*_b_*, in LMM with *X* as matrix of fixed effects, *ZL*^−1^ as matrix of random effects, *b* ~ 𝒩(0, σ^2^*_b_I_p_*) and *ε* ~ 𝒩(0, σ^2^*Ι_η_*);

**4:** define λ := σ̂^2^/σ̂^2^*_b_* and find estimates *β̂*^crP^ and *b̂*^crP^ by applying formula (2.4) with matrix *Q̃*.

The value of λ_2_ is fixed as a small number. In our work, we tested several values of λ_2_ to assess the solutions’ stability. Based on the simulation results, we fix λ_2_ to be equal to 0.001.

### 3.2 riPEER

In our second approach, we propose to use two penalty parameters in the optimization problem (3.1) – main penalty parameter λ_*Q*_ and a ridge adjustment parameter λ_2_. Solution to this problem is found using the equivalent LMM formulation estimating both penalty parameters. Specifically, the optimization problem can be written as,

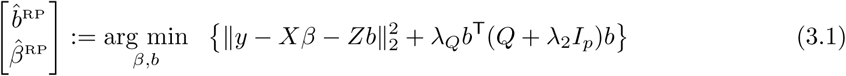

or equivalently as

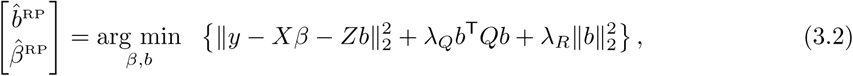

where λ_*R*_ = λ_*Q*_λ_2_. The latter formulation, with an additional ridge penalty, justifies the name of the method. Without loss of generality, in the first step we can reduce the problem by excluding the Χ matrix. It can be shown that

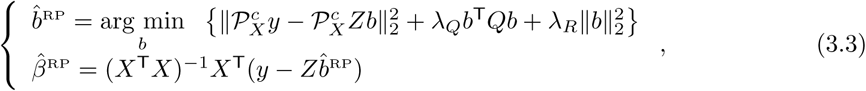

where 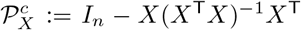 is the projection onto orthogonal complement to the subspace spanned by columns of *X*. Denote 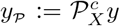 and 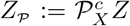. For the ease of computation, we perform an additional step; let *USU*^T^ be the SVD of the matrix *Q*, with the orthogonal matrix *U* and the diagonal matrix *S* with nonnegative numbers on the diagonal. After the variable change via the transformation *b* ← *U*^T^*b*, the final form of problem becomes

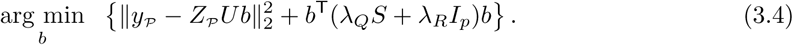

Now the LMM equivalent to (3.4) can be characterized as follows:

B.1 We consider model *y_P_* = *Z̃b* + *ε*, with *Z̃* := *Z_P_U* and *b* being the vector of random effects,
B.2 *ε* ~𝒩 (0,σ^2^*Ι_n_*),
B.3 *b* ~ 𝒩(0,σ^2^D^−1^_λ_), where *D*_λ_ := λ*_Q_S* + λ*_R_Ι_p_*.

To the best of our knowledge, there is no existing *R* package that can be used to estimate the parameters based on the equation (3.4) in a straightforward way. Thus, we have focused on deriving and optimizing the LMM log-likelihood function. Following Demidenko (2004), maximizing the log-likelihood is equivalent to minimizing the expression

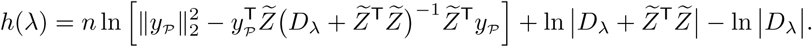

Precisely, the maximum likelihood estimator (**mle**) for σ^2^, λ*Q* and λ_*R*_ can be found as

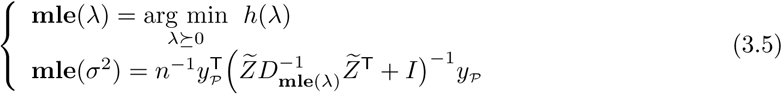

To find the solution to the above optimization problem we utilized *sbplx* function from the *nloptr R* software package (Ypma, 2014), which is an *R* interface to NLopt (Johnson, 2016) – a free/open-source library for nonlinear optimization. *sbplx* function implements Subplex algorithm (Rowan, 1990) to estimate the parameters of an objective function to be minimized. riPEER algorithm can be summarized as

#### Algorithm 2 riPEER

**input:** matrices *Z, X*, positive semi-definite matrix *Q*

**1:** find SVD of *Q*, i.e. *Q* = *USU*^T^;

**2:** define projection matrix 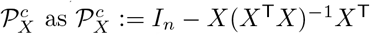

**3:** denote 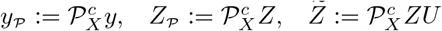

**4:** find estimates of λ_*Q*_ and λ_*R*_ by solving optimization problem in (3.5);

**5:** define 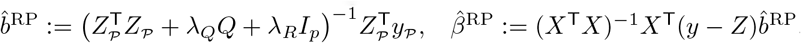.

### 3.3 Variable Reduction PEER (vrPEER)

For the sake of considering an alternative approach that does not introduce an additional penalty term to deal with the singularity of a matrix *Q*, i.e. rank(*Q*) < *p,* we reduce the number of penalized predictors in the optimization problem by moving a number of them to the unpenalized matrix *X*. This method is based on the property stating that for any vector *b* ∈ ℝ^*p*^, *b*^T^*Qb* can be expressed as *c*^T^ *diag*(*s_1_,…, s_r_*)*c* for nonzero singular values *s*_1_,…, *s_r_* of *Q* and some c from the *r*-dimensional subspace spanned by the singular vectors corresponding to *s*_1_,…, *s_r_*.

In the next step, we proceed with the variable rearrangement. We start with the optimization problem (2.2) with *Q* having rank *r* < *p*. Matrix *Q* has an eigendecomposition of the form

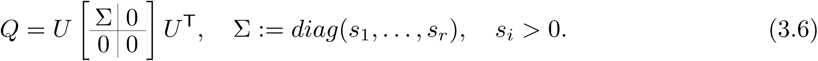

Using the notation *b̃* := *U*^T^*b* and *b̃* _[1:*r*]_ for the first *r* coefficients of *b̃*, we get the equivalent optimization problem with the objective function

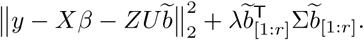

Matrix *ZU* is separated column-wise into A which is a submatrix of *ZU* created by the last *p* − *r* columns and ℤ which is a submatrix of *ZU* created by the first *r* columns. Let *c* := *b̃* [_*r*+1*:p*_] be the regression coefficients corresponding to the columns in matrix *A* and *d* := *b̃*_[1:*r*]_ the regression coefficients corresponding to the new penalized variables in the matrix ℤ. Now, the optimization criterion can be written as,

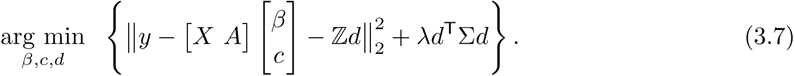

Optimization problem (3.7) is in the ordinary PEER optimization form (2.2) with an invertible penalty matrix Σ. Thus, an equivalent LMM can be easily derived. In summary, for a singular matrix *Q* some directions, corresponding to the singular vectors with zero singular values in ℝ^*p*^, are not penalized. After changing the basis vectors, we can move the corresponding variables to the unpenalized part. All the steps are summarized in the Algorithm 2.

#### Algorithm 3 vrPEER

**Input:** matrices *Z, X,* positive semi-definite matrix *Q* with rank *r* < *p*;

**1:** Let *U* be orthogonal matrix from (3.6) and define matrix of nonzero singular values, Σ :=*diag*(*s*_1_, …, *s_r_*);

**2:** Denote by *A* last *p – r* columns of *ZU* and by ℤ first *r* columns of *ZU*;

**3:** Put 𝕏 := [*X A*] and consider minimization problem with the objective 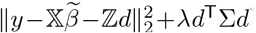, for *β̃* ∈ ℝ^*m*+*p*−*r*^ and *d* ∈ ℝ^*r*^

**4:** Estimate σ^2^ and *σ*^2^_d_ in equivalent LMM, i.e. the model *y* = 𝕏**β̃**+ℤ*d*+*ε*, where 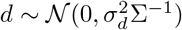, and *ε* ~ 𝒩(0, *σ*^2^*I_n_*), and define 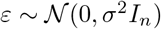;

**5:** Find estimates *β̂*;^vrΡ^ and *b̂* ^vrP^ by applying formula (2.4)

In Section 4, we study the empirical performance of the proposed extensions to PEER. Namely, we conduct an extensive simulation study using data-driven connectivity matrices to evaluate the estimation performance of riPEER-c, vrPEER and riPEER. In addition, we compare the results from our proposed methods with the ordinary ridge regression and ordinary least squares.

## 4 Simulation experiments

In this section, we study the empirical behavior of the extensions to the PEER-type estimators proposed, namely riPEER-c, riPEER and vrPEER, alongside the ordinary ridge and PEER estimation methods in a number of simulation studies.

In Section 4.1, we summarize the setup of our simulation studies. We concentrated on the cases with both full and partial information provided by the connectivity matrices. In Section 4.2, methods used for the parameter estimation are discussed. Finally, in Section 4.3, we summarize the estimation discrepancies for the proposed and classical methods including the comparison between two approaches to penalty parameter(s) choice: cross-validation technique and REML estimation within the linear mixed models framework.

### 4.1 Simulation scenarios

We aim to evaluate the methods' behavior assuming that connectivity information among the variables is at least partially informative. In practice, we presume the existence of some a priori knowledge about the structural or functional connectivity among the cortical brain regions (see Section 5). Here, we investigate scenarios in which connectivity information accurately reflects the structure of variables as well as when it does not, starting from a low fraction of connections being permuted to a completely uninformative setting.

In each simulation scenario, we first generate a matrix of p correlated features, *Z* ∈ ℝ^*n*×*p*^, where the rows are independently distributed as 𝒩_*p*_(0, Σ) with ∑_*ij*_ = *exp* {−*k*(*i* − *j*)^2^} , *i* = 1,… ,*p*; *j* = 1,… ,*p*. We then generate a vector of true coefficients *b* ~ 𝒩(0,σ^2^_*b*_*Q*^−1^), reflecting the underlying connectivity information. Here, *Q* denotes the normalized Laplacian of an adjacency matrix 𝒜 that represents the assumed connectivity between the variables; see Section 2.2. All simulation scenarios are constructed by perturbing an established modularity matrix obtained by Sporns (2013); see also (Cole et al., 2014; Sporns and Betzel, 2016) in which each node belongs to one of five modules (blocks). This base adjacency matrix was produced in the FreeSurfer software (Fischl, 2012) and is displayed in Figure 10 in the Appendix.

Our simulation are conducted under a wide range of scenarios involving various choices of simulation settings:

1. number of predictors: *p* ∈ (100, 200)
2. number of observations: *n* ∈ (100, 200)
3. signal strength: σ_*b*_^2^ ∈ (0.1, 0.01)
4. strength of correlation between the variables in the *Z* matrix: *k* ∈ (0.01, 0.004)
5. information content in the adjacency matrix 𝒜: complete information or partial information

All of these settings effect the estimation accuracy of the coefficient vector, *b* ∈ *R^p^,* representing the association between the *p* variables in *Z* and outcome *y* in the model *y* = *Zb* + *ε*, where *ε* ~ 𝒩 (0,σ^2^*I_n_*). To evaluate the performance of various estimation procedures, we must define a coefficient vector, *b*, whose structure reflects that of adjacency matrix 𝒜. This is done by generating 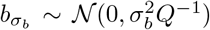 for various values of σ^2^_*b*_, which may be viewed as signal strength for a fixed value of error variance σ^2^ = 1.

In the simulations, we consider two scenarios regarding the information provided by *Q*: (1) complete knowledge of connectivity and (2) misspecified connectivity. In the former, the normalized Laplacian *Q* is constructed using the “true” adjacency matrix 𝒜, while in the latter, *Q* is constructed from a perturbed adjacency matrix 𝒜^*obs*^ of the “true” 𝒜. To implement varying amounts of “true” information in 𝒜^*obs*^, we vary parameters, *steps* ∈ {0,10,100,1000,10000} and *prob* ∈ {0,0.2,0.4,0.6,0.8,1}, that effectively “rewire” the adjacency matrix 𝒜 (see details in a paragraph on page 11). We denote these perturbed matrices by 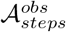 and 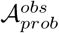, respectively.

For each set of simulation scenarios, *Z_n,p_*, 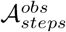 or 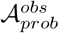 and 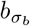, we perform 100 experimental runs and for each run we simulate the outcome via 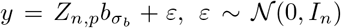. Finally, we assess the performance of all considered methods by comparing relative mean squared error (MSE) of the estimated coefficients *b* via 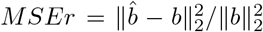, where ||*x*||_2_ denotes the *l*^2^-norm of a vector *x* = (*x*_1_,…, *x_p_*), given by 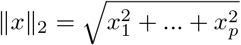. Motivated by the applications to the brain imaging data, we focus on the accuracy of *b* estimation.

**Figure 2:**
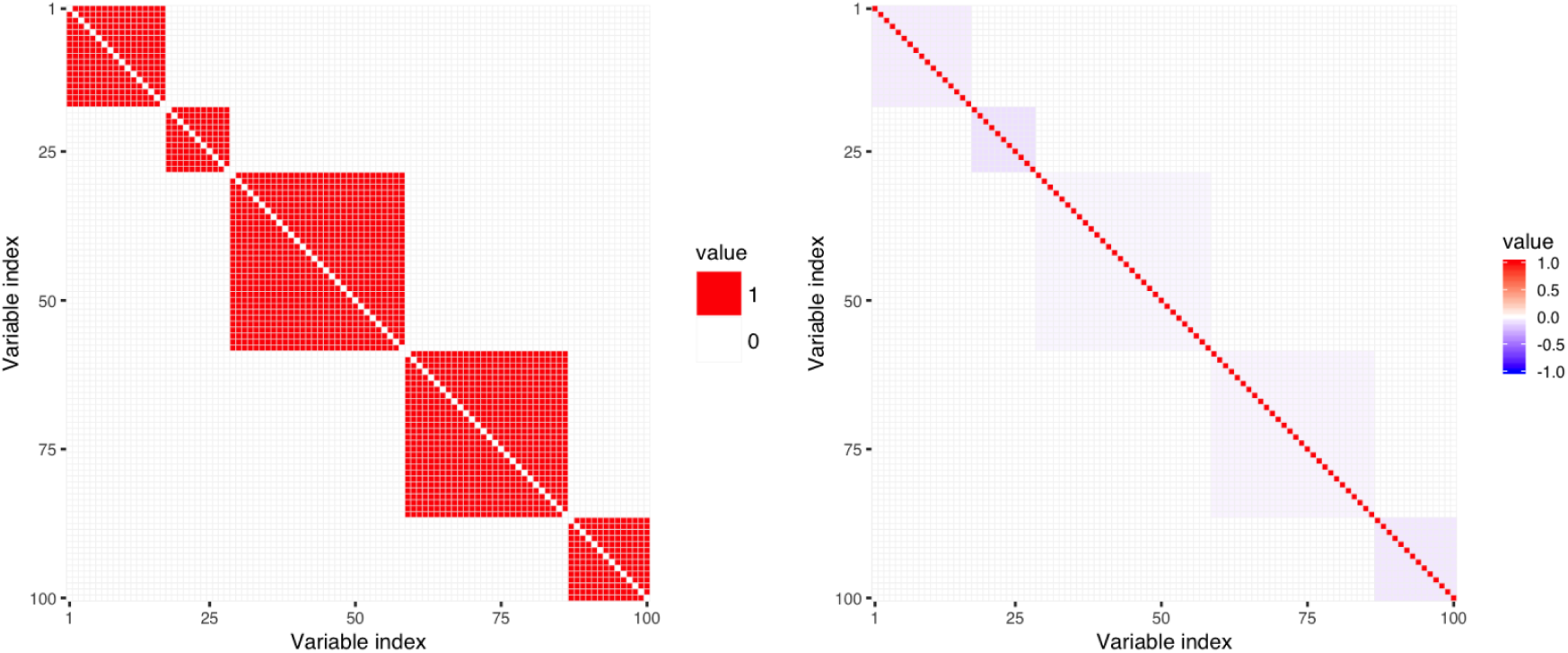
64 × 64 dimensional *modularity* adjacency matrix is rescaled to 100 × 100 dimensional adjacency matrix (left panel). Corresponding normalized Laplacian matrix is shown in the right panel.

**Figure 3:**
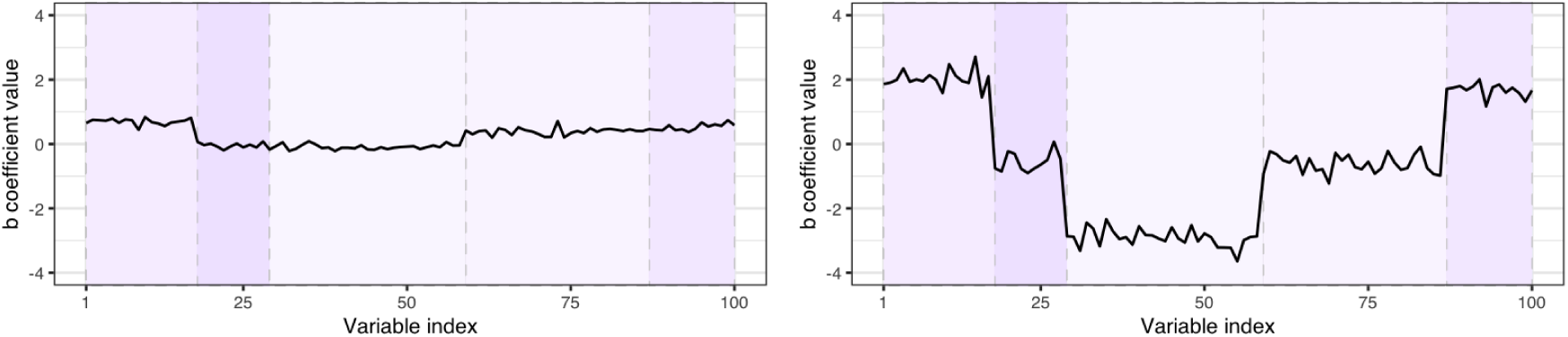
Exemplary *b* values simulated from *b* ~ 𝒩(0,σ^2^_*b*_*Q−1*) distribution for σ^2^_*b*_ (left panel) and σ^2^_*b*_ (right panel), for *Q* being the Laplacian matrix of 100 × 100 dimensional graph adjacency matrix obtained by rescaling original *modularity* graph adjacency matrix. Colors of the background rectangular shades correspond to the color scale of the Laplacian matrix *modules* in Figure 2 (right panel).

#### Construction of the graph adjacency matrix 𝒜^*obs*^

To express misspecified connectivity information in the simulation setting, we construct the normalized Laplacian matrices based on either 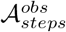 or 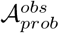 which are permuted versions of the true connectivity matrix 𝒜. Entries of 𝒜 are permuted (“rewired”) using one of the following methods:

1. 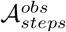 is generated via an iterative randomization procedure which preserves the original graph’s density, degree distribution and degree sequence, as implemented by keeping_degseq function from igraph R package (Csardi and Nepusz, 2006). The rewiring algorithm chooses two arbitrary edges in each step, (*a, b*) and (*c, d*), and substitutes them with (*a, d*) and (*c, b*), if they not already exists in the graph. Function’s *niter* argument denotes the number of *steps* (rewiring trials) performed.
2. 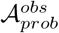 is generated via a randomization procedure which preserves the original graph’s density, as implemented by each_edge function from igraph R package. The function rewires the endpoints of the edges with a constant probability uniformly randomly to a new vertex in a graph. Function’s *prob* argument denotes a constant probability for a graph node to be rewired.

Simulation parameters steps and *prob* are therefore reflecting the level of inaccuracy in 𝒜^*obs*^ matrix obtained with the use of the first and second method, respectively, with higher parameter values indicating higher inaccuracy of the connectivity information. 𝒜^*obs*^ examples are presented in Figures 4 and 5.

**Figure 4:**
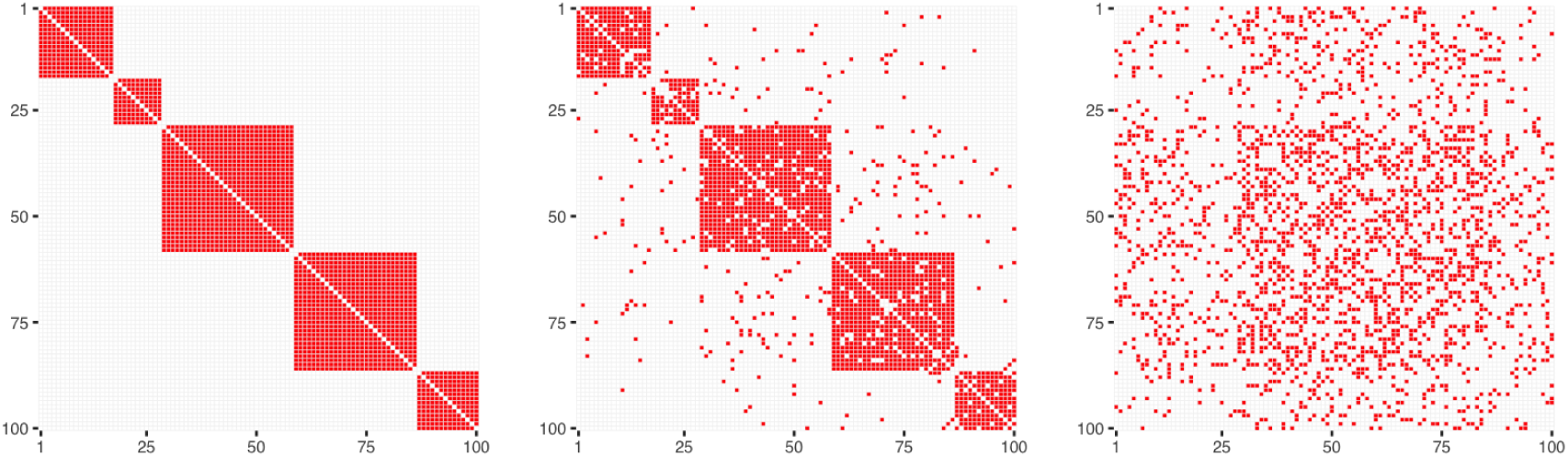
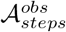 matrices obtained from a graph rewiring while preserving the original graph's degree distribution, for *p* = 100 and *steps* = 0 (left panel), *steps* = 100 (central panel), *steps* = 10000 (right panel).

**Figure 5:**
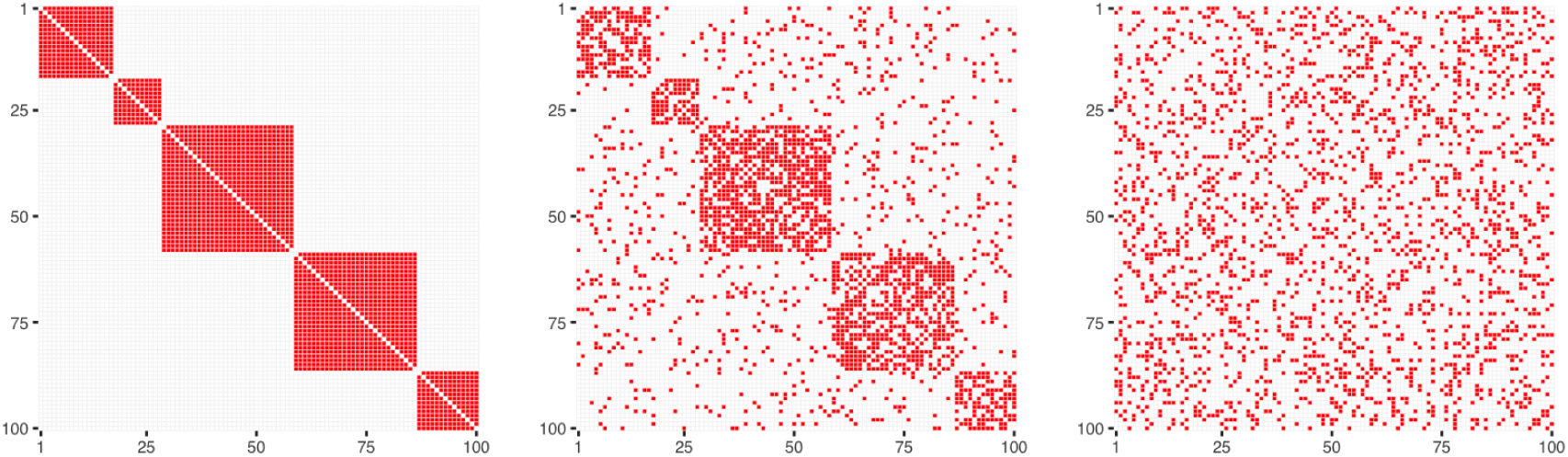
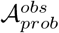 matrices obtained from a graph rewiring with a constant probability for a graph node to be rewired, for *p* = 100 and *prob* = 0 (left panel), *prob* = 0.2 (central panel), *prob* = 1 (right panel).

The two rewiring methods have different properties. Rewiring while preserving the original graph’s degree distribution yields 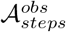 matrix that still reflects the initial *modules* size proportions even for extremely high *steps* parameter values (Figure 4, right panel), while rewiring with a constant probability for a graph node to be rewired yields completely random distribution of connections across a graph (Figure 5, right panel).

### 4.2 Parameter estimation

In the simulation study, we compare the following methods of fitting a linear regression model *y* = *Zb* + *ε*: ordinary ridge using two approaches to penalty parameter choice — CV and REML, PEER – CV, riPEER-c — REML, vrPEER — REML, and riPEER — CV and REML.

In the simulation study, we utilize mdpeer R package (Karas, 2016) for the parameter estimation for all the methods, except for an ordinary ridge with λ_*R*_ regularization parameter chosen by cross-validation, which uses the implementation from glmnet R package (Friedman et al., 2010). For the latter, we performed 10-fold cross-validation (CV) where a loss function is a squared-error of response variable’s predictions. We used the default setting from the `glmnet::coef.glmnet` which selects λ_*R*_ to be the largest value of λ such that the CV error is within 1 standard error of the minimum CV. For the PEER method the regularization parameter λQ was chosen by 10-fold cross-validated search over a grid: 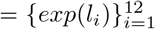, for 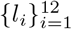 being a sequence of equally spaced values between –4 and 9. The selected λ_Q_ value yields the minimum squared prediction error. riPEER regularization parameters (λ*_Q_*, λ*_R_*) were chosen by 10-fold cross-validation searching over a fixed 2dimensional grid of (λ*_Q_*, λ*_R_*) parameter values: 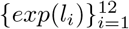, where 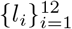 is a sequence of equally spaced values between –4 and 9. We selected a pair of values (λ*_Q_*, λ*_R_*) which yield the minimum squared prediction errors averaged over the 10 folds.

### 4.3 Simulation results

We investigated the estimation performance of all methods in the cases with both full and partial information provided by the connectivity matrices. For the simulation experiments with full information, we considered methods which use both CV and REML to estimate the penalty parameter(s) (4.3.1). For the simulation experiments with partial information provided, we narrowed down our investigation to the methods which use REML estimation for the regularization parameter choice and considered simulation scenarios in which entries of 𝒜 from the true *b* simulation are permuted with the use of two different methods (4.3.2 and 4.3.3).

#### 4.3.1 Informative connectivity information input

Table 1 summarizes the relative mean squared error of estimation 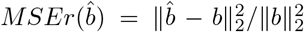 averaged over 100 simulation runs for each simulation scenario. Minimum *MSEr* values are highlighted (bolded) for each row. It is important to note that since the true regression vector 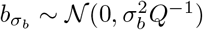 was generated for each simulation scenario separately, the results are comparable within rows, but not across them.

One can observe that methods that use REML outperform the corresponding methods using CV for the selection of the tuning parameter(s). More precisely, *MSEr*(*b̂*) is lower by 17.9% on average for REML than for CV in the ridge penalty methods group, by 12.7% in the PEER penalty methods group and by 13.0% in the riPEER penalty methods group, respectively. A close examination reveals the presence of simulation “outliers” causing larger *MSEr*(*b̂*) for PEER–CV and riper–CV as opposed to riPEER-c/vrPEER-REML and riPEER–REML, respectively (see Figures 12 and 13 in the Appendix A). This is consistent with the work of P. T. Reiss (2009), where the authors noted that in their simulations cross-validation failed to find the global optimum of corresponding optimization problem more often than REML.

In addition, we observe major advantages of the methods using PEER-type penalty term over a method using ordinary ridge penalty term. In particular, vrPEER–REML yields lower values of *MSEr* error than ridge–REML for each combination of the simulation parameters. It is on average lower by 69.5%. Also, we note that riPEER-c and vrPEER methods yield almost identical results.

Finally, the proposed riPEER–REML method behaves as anticipated; it mirrors the results of a method with a penalty term exhibiting better *b* estimation performance in particular simulation setting. Since the connectivity assumed in these simulation settings is fully informative, riPEER–REML approximates the results of the graph-constrained regression methods riPEER-c/vrPEER–REML rather than ridge–REML which does not utilize connectivity matrix in the estimation process.

#### 4.3.2 Partially informative connectivity matrix input: graph rewiring with the original graph’s degree distribution preserved – 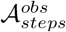

In the simulation experiments, we studied 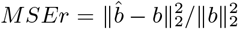 changes when the original connectivity information graph is rewired and used for the regularized estimation of *b*. In this experiment, we used a graph rewiring method that preserves the original graph’s degree distribution. Figure 6 summarizes *MSEr* averaged over 100 simulation runs for *k* = 0.01, σ^2^_*b*_, *p* ∈ {100, 200} (left and right panels, respectively) and *n* ∈ {100,200} (top and bottom panels, respectively), *steps* ∈ {0,10,100,1000,10000} (marked by x-axis labels), for REML estimation methods: ridge, riPEER-c, vrPEER, riPEER.

**Figure 6:**
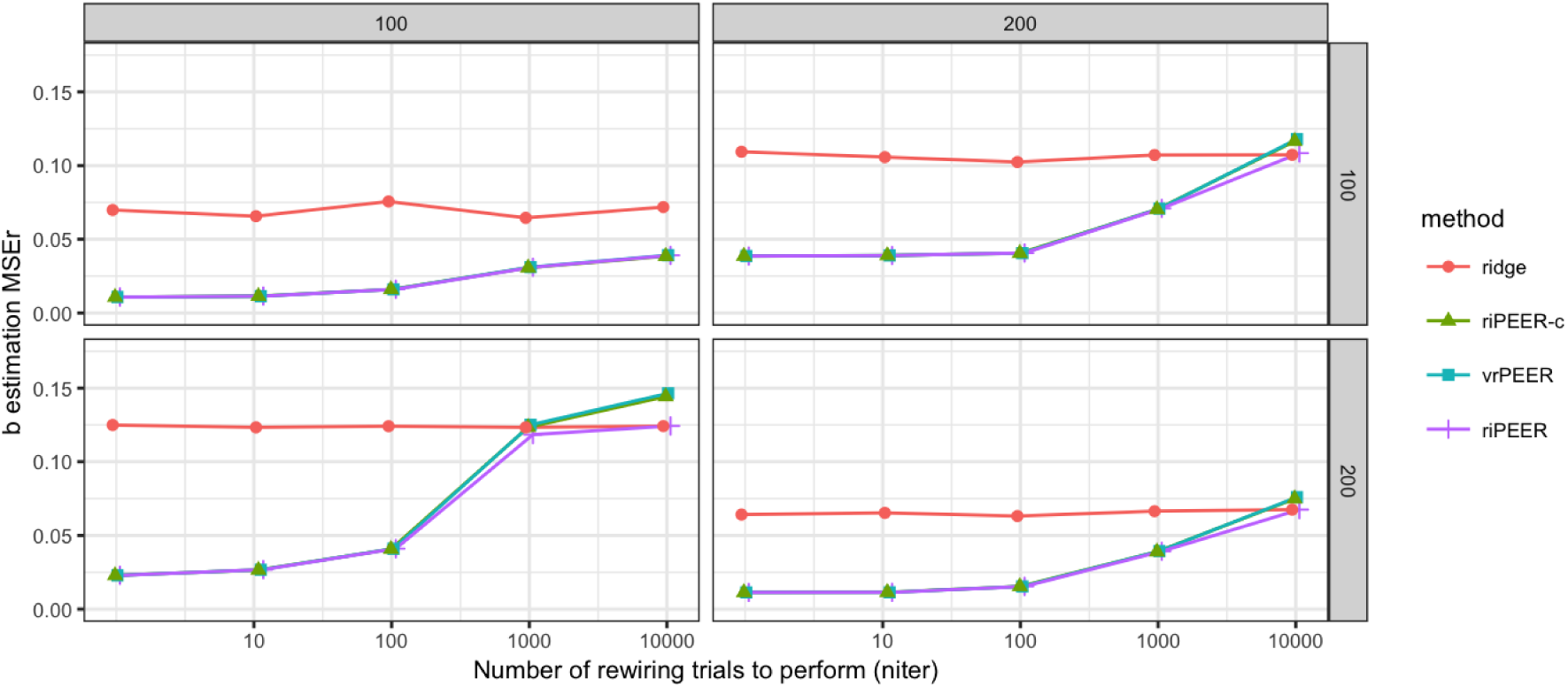
*b* estimation relative error 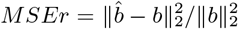 values for *k =* 0.01, σ^2^_*b*_ = 0.01, *p* ∈ {100, 200} (left and right panels, respectively) and n ∈ {100, 200} (top and bottom panels, respectively), *steps* ∈ {0,10,100,1000,10000} (marked by x-axis labels), for REML estimation methods: ridge, riPEER-c, vr-PEER, riPEER, in case of the uninformative connectivity information input (graph rewiring with the original graph's degree distribution preserved).

Figure 6 displays the summary of the relative estimation errors for all methods. When connectivity information is exploited in estimation, riPEER-c, vrPEER and riPEER outperform or behave no worse than the ridge estimation method in simulation scenarios with low and moderate amount of original connectivity information graph rewiring (*steps* ∈ {0,10,100,1000}). For the extreme amounts of original connectivity information graph rewiring (*steps* = 10000), riPEER-c and vrPEER start to perform slightly worse than ridge. riPEER exhibits property of adaptiveness towards the amount of true information contained in the penalty matrix; it yields results like riPEER-c or vrPEER estimators in simulation scenarios with low and moderate amount of original connectivity information graph rewiring and yields results similar to ridge for extreme amount of original connectivity information graph rewiring. Therefore, riPEER performs no worse than any other method considered, regardless of the amount of true information contained in the penalty matrix. Notably, riPEER-c and vrPEER yield almost identical results for each combination of simulation parameters considered in this experiment.

Similar trends of estimation methods performance are observed for other combinations of simulation parameter values – a complete set of *MSEr* values obtained is presented in Figures 14 and 15 (Appendix A) and summarized in Table 3 (Appendix A).

**Table 1:** *b* estimation relative error 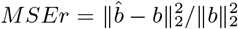 averaged over 100 simulation runs for each combination of the simulation parameters and each method, in case of informative connectivity information input. Minimum estimation *MSEr* values are highlighted (bolded) for each row.

| $n$ | $p$ | $k$ | $\sigma_b^2$ | ridge<br>CV | ridge<br>REML | PEER<br>CV | riPEER-c<br>REML | vrPEER<br>REML | riPEER<br>CV | riPEER<br>REML |
| --- | --- | --- | --- | --- | --- | --- | --- | --- | --- | --- |
| 100 | 100 | 0.004 | 0.01 | 0.341 | 0.133 | **0.015** | **0.015** | **0.015** | 0.023 | **0.015** |
| 100 | 100 | 0.004 | 0.1 | 0.220 | 0.096 | **0.017** | **0.017** | **0.017** | 0.019 | **0.017** |
| 100 | 100 | 0.01 | 0.01 | 0.309 | 0.170 | 0.065 | **0.030** | **0.030** | 0.032 | **0.030** |
| 100 | 100 | 0.01 | 0.1 | 0.136 | 0.067 | 0.012 | **0.011** | **0.011** | 0.013 | **0.011** |
| 100 | 200 | 0.004 | 0.01 | 0.157 | 0.123 | 0.031 | **0.030** | 0.031 | 0.033 | 0.031 |
| 100 | 200 | 0.004 | 0.1 | 0.238 | 0.128 | 0.034 | **0.031** | **0.031** | 0.037 | **0.031** |
| 100 | 200 | 0.01 | 0.01 | 0.197 | 0.164 | 0.091 | **0.083** | **0.083** | 0.087 | **0.083** |
| 100 | 200 | 0.01 | 0.1 | 0.326 | 0.169 | 0.114 | **0.100** | **0.100** | 0.107 | **0.100** |
| 200 | 100 | 0.004 | 0.01 | 0.078 | 0.128 | 0.092 | **0.042** | **0.042** | 0.056 | **0.042** |
| 200 | 100 | 0.004 | 0.1 | 0.093 | 0.087 | **0.012** | **0.012** | **0.012** | 0.014 | **0.012** |
| 200 | 100 | 0.01 | 0.01 | 0.077 | 0.130 | 0.049 | **0.038** | 0.039 | 0.044 | 0.039 |
| 200 | 100 | 0.01 | 0.1 | 0.129 | 0.166 | 0.072 | **0.068** | **0.068** | 0.076 | **0.068** |
| 200 | 200 | 0.004 | 0.01 | 0.252 | 0.213 | 0.104 | **0.102** | **0.102** | 0.108 | **0.102** |
| 200 | 200 | 0.004 | 0.1 | 0.129 | 0.077 | 0.023 | **0.019** | **0.019** | 0.025 | **0.019** |
| 200 | 200 | 0.01 | 0.01 | 0.117 | 0.101 | 0.030 | **0.029** | **0.029** | 0.031 | **0.029** |
| 200 | 200 | 0.01 | 0.1 | 0.166 | 0.123 | 0.063 | **0.058** | **0.058** | 0.065 | **0.058** |

#### 4.3.3 Partially informative connectivity matrix input: graph rewiring with a constant probability for a graph node to be rewired 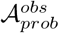

In this experiment with partially informative connectivity input we used a graph rewiring method with a constant probability for a graph node to be rewired. Therefore, the initial *modules'* size proportions of a graph connectivity matrix are no longer apparent for high *prob* parameter values.

Figure 7 summarizes *MSEr* averaged over 100 simulation runs for *k* = 0.01, σ^2^_*b*_, *p* ∈ {100, 200} (left and right panels, respectively) and *n* ∈ {100, 200} (top and bottom panels, respectively), *prob* ∈ {0, 0.2,0.4, 0.6,0.8,1} (marked by x-axis labels), for REML estimation methods: ridge, riPEER-c, vrPEER, riPEER. Similarly to the previous experiment, we can observe that methods which utilize connectivity information as *a priori* knowledge in the estimation outperform the ordinary ridge estimation method in simulation scenarios with low and moderate amount of original connectivity information graph rewiring (*prob* ∈ {0, 0.2,0.4}). Also, the adaptive behavior of riPEER is apparent as it approximates results of either ridge or riPEER-c/vrPEER, depending on which of these methods yields lower *MSEr* value in a particular simulation setting. Again, riPEER-c and vrPEER yield almost identical results for each combination of the simulation parameters considered in this experiment.

**Figure 7:**
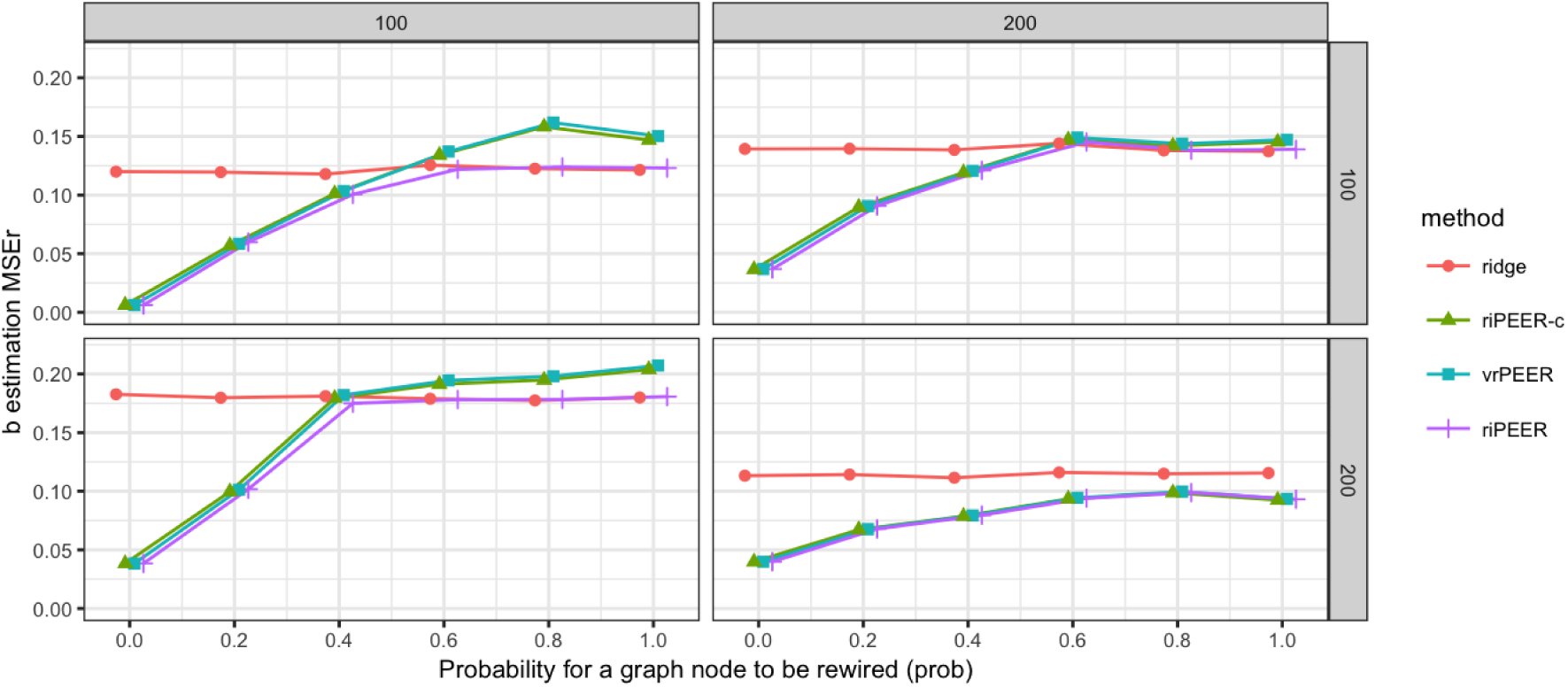
*b* estimation relative error 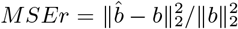 values for *k* = 0.01, σ^2^_*b*_, *p* ∈ {100, 200} (left-side and right-side panels, respectively) and *n* ∈ {100, 200} (upper-side and bottom-side panels, respectively), *prob ∈* {0, 0.2, 0.4, 0.6, 0.8,1} (marked by x-axis labels), for “REML” estimation methods: ridge, riPEER-c, vrPEER, riPEER, in case of the uninformative connectivity information input (graph rewiring with a constant probability for a graph node to be rewired).

Similar trends of estimation methods performance are observed for other simulation parameter values – a complete set of *MSEr* values is vizualized in Figures 16 and 17 (Appendix A) and summarized in Table 4 (Appendix A).

The numerical experiments conducted show that with an informative connectivity information input, PEER-based regularization methods yield lower average *MSEr* values than the ordinary ridge method. In addition, we show that the proposed riPEER method, which combines ordinary ridge and PEER-based penalty, is adaptive to the level of information present in a connectivity matrix. This property stems from the data-driven estimation of the regularization parameters for the ordinary ridge and PEER penalty terms. More precisely, riPEER estimator’s performance is similar to a PEER estimator when the connectivity matrix is informative, whereas for an uninformative connectivity matrix riPEER behaves like ordinary ridge.

In addition, we observe that REML, compared to CV, provides estimates resulting in a smaller *MSEr* for each simulation setting and within each estimation method studied. This is consistent with the observations of (P. T. Reiss (2009)).

## 5 Imaging data application

Using the proposed methods, we model the associations between alcohol abuse phenotypes and structural cortical brain imaging data. Specifically, we utilize cortical thickness measurements obtained using the FreeSurfer software (Fischl, 2012) to predict alcoholism-related phenotypes while incorporating the structural connectivity between the cortical regions obtained by Sporns (2013); see also Cole et al. (2014); Sporns and Betzel (2016).

Our results for the cortical thickness association were obtained in a large sample of young, largely nonsmoking drinkers. This male-only alcoholism risk sample is very different from the studies of older alcoholic subjects recently reported in several articles (Momenan et al., 2012; Nakamura-Palacios et al., 2014; Pennington et al., 2015). The number of reported morphometric differences found tended to involve the frontal lobe, with the effects considerably weaker in men only and after accounting for substance use. There is no consensus about the laterality of the cortical thinning effects in alcoholics, albeit some of the findings are medial (anterior cingulate cortex, ACC), while others could be explained by more prominent right hemisphere effects of lifetime alcohol exposure (Momenan et al., 2012) not yet evident in younger population.

### 5.1 Data and preprocessing

#### Study sample

The sample consisted of 148 young (21-35 years) social-to-heavy drinking male subjects from several alcoholism risk studies. Structural imaging data from 88 subjects were included in the study relating externalizing personality traits and gray matter volume (Charpentier et al., 2016), with a subset of these 88 subjects also reported in the studies of dopaminergic responses to beer (Oberlin et al., 2013, 2015, n=49 and 28, respectively), and regional cerebral blood flow (Weafer et al., 2015, n=44). Subjects’ demographic and a risk for alcoholism-related characteristics are summarized in Table 2.

**Table 2:** Study sample subjects’ characteristics. ^*a*^Number of first or second degree relatives with Alcohol Use Disorder by self-report. ^*b*^Alcohol Use Disorders Identification Test. ^*c*^Drinking data are from the Timeline Followback Interview. ^*d*^ Drinking frequency data from past 35 days of a subject’s history.

| Study variables | Min | Max | Mean | Median | StdDev |
| --- | --- | --- | --- | --- | --- |
| Age | 21.0 | 35.0 | 24.0 | 23.0 | 3.2 |
| Education | 11.0 | 20.0 | 15.2 | 15.0 | 1.5 |
| AUD relatives<sup>a</sup> | 0.0 | 7.0 | 1.2 | 1.0 | 1.5 |
| AUDIT<sup>b</sup> | 3.0 | 28.0 | 10.9 | 10.0 | 4.9 |
| Smoker | 0.0 | 1.0 | 0.1 | 0.0 | 0.2 |
| Drinks per week<sup>c,d</sup> | 0.4 | 62.0 | 16.3 | 14.9 | 10.6 |
| Drinks per drinking day<sup>c,d</sup> | 1.3 | 23.9 | 5.9 | 5.5 | 3.4 |
| Heavy drinking days per week<sup>c,d</sup> | 0.0 | 4.4 | 1.4 | 1.4 | 1.1 |
| Years since first drink<sup>c</sup> | 2.0 | 24.0 | 7.7 | 7.0 | 4.1 |
| Years since regular drinking<sup>c</sup> | 0.0 | 17.0 | 5.2 | 4.0 | 3.6 |
| Years since first intoxication<sup>c</sup> | 2.0 | 24.0 | 6.9 | 6.0 | 3.9 |

#### MRI data acquisition

Brain imaging was performed using the following Siemens 3T MRI scanners/head coils: Trio-Tim/12-channel, Skyra/20-channel and Prisma/20&64-channel at the Indiana University School of Medicine Center for Neuroimaging. For all configurations, whole-brain high resolution anatomical MRI was collected using a 3D Magnetization Prepared Rapid Acquisition Gradient Echo (MP-RAGE) with imaging parameters optimized according to the ADNI (Alzheimer’s Disease Neuroimaging Initiative) protocol (Prisma/Skyra, 5.12 min, 1.05 × 1.05 × 1.2 mm^3^; Trio-TIM; 9.14 min, 1.0 × 1.0 × 1.2 mm^3^ voxels).

#### Cortical measurements

The FreeSurfer software package (version 5.3) was used to process the acquired structural MRI data, including gray-white matter segmentation, reconstruction of cortical surface models, labeling of regions on the cortical surface and analysis of group morphometry differences. The resulting dataset has cortical measurements for 68 cortical regions with parcellation based on Desikan-Killiany atlas (Desikan et al., 2006) was computed. The subset of 66 variables describing average gray matter thickness (in millimeters) of gray matter brain regions did not incorporate left and right insula due to their exclusion from the structural connectivity matrix. It is important to note that the cortical measurement variables exhibit multicollinearity as evident in the high values of pairwise Pearson correlation coefficient (Figure 18 in the Appendix A).

#### Structural connectivity information

Structural connectivity was estimated based on a model proposed by Hagmann et al. (2008) using a diffusion map to construct 3D curves of the maximal diffusion coherence. This model estimated a structure of five connectivity modules. Binary information of a common assignment to one of the modules was identified for each pair of the 66 cortical regions. The resulting 66 × 66 matrix 𝒜 was used to represent similarities between all brain region pairs. This matrix, illustrated in Figure 10 in the Appendix A, played the role of a structural connectivity adjacency matrix that was used to define *Q*, illustrated in Figure 11 in the Appendix A.

### 5.2 Estimation methods

We employed ordinary ridge, riPEER-c, riPEER and vrPEER with REML estimation of the regularization parameter(s) to quantify the association of imaging markers with the drinking frequency. In addition, we fitted both simple and multiple linear regression models to compare the unpenalized regression estimates to the estimates obtained from the regularized regression approach. We used the derived cortical thickness measures from 66 brain regions as predictors of the outcome *y* = *Number of drinks per drinking day* and connectivity matrix-derived Laplacian was used as a penalty matrix for riPEER-c, riPEER and vrPEER. For each of the four regularized regression estimation methods, we included *Age, Smoker* and *Years since the start of regular drinking* as non-penalized adjustment variables. Both outcome and predictors were standardized to mean zero and variance equal to one.

### 5.3 Results

The regression coefficient estimates obtained from both simple and multiple linear regression are presented as black solid lines in Figure 8 in the top and bottom panel, respectively. Statistically significant coefficients (at a nominal level equal to 0.05) are marked with solid red vertical lines.

**Figure 8:**
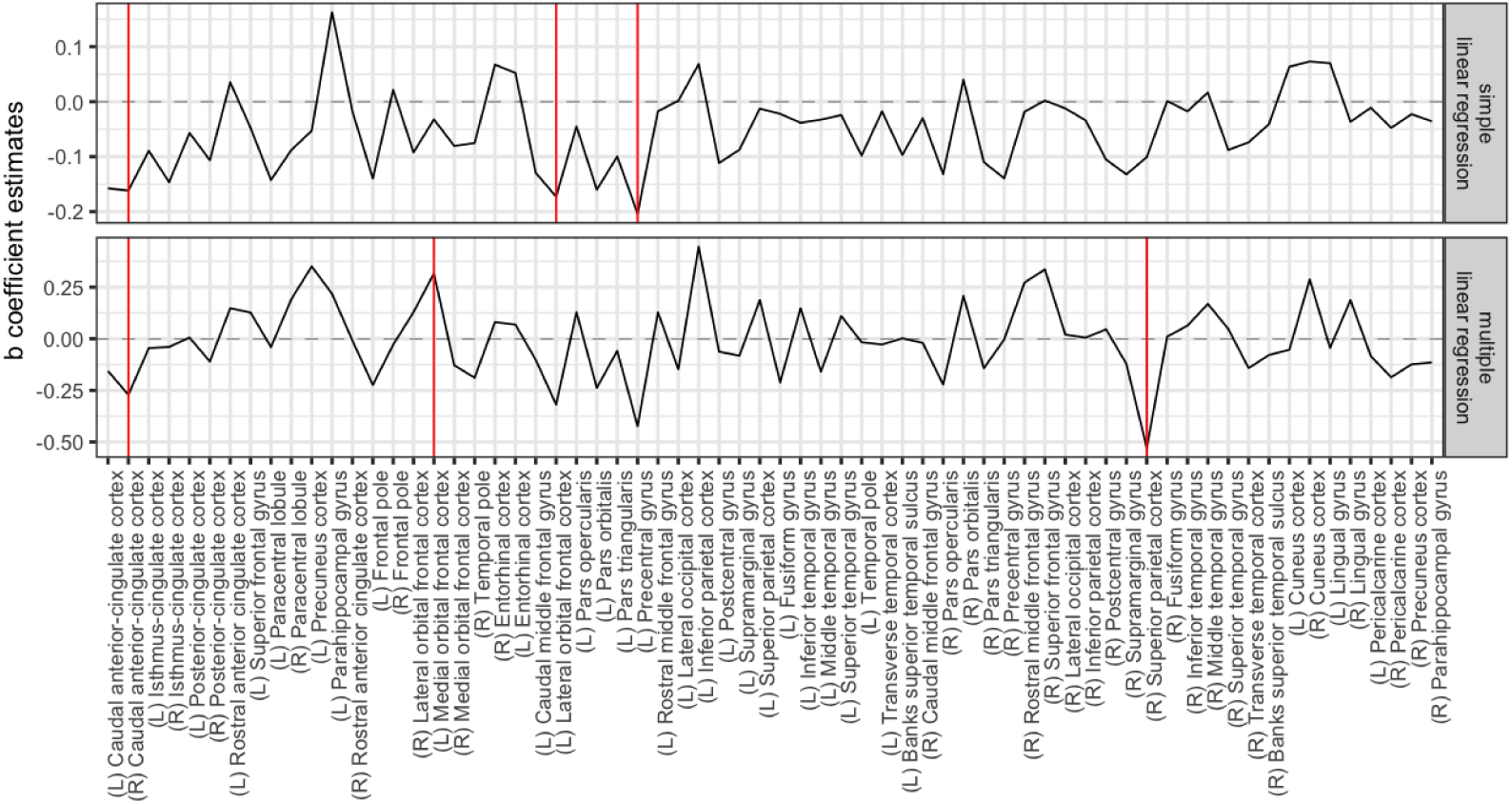
*b̂* coefficient estimates obtained from linear regression: simple (upper panel) and multiple (bottom panel). Estimates’ values are denoted with a solid black line. Statistically significant (with a significance level of 0.05) coefficients are denoted with solid red vertical lines.

The regression coefficient estimates obtained from the ordinary ridge, riPEER-c, riPEER and vrPEER are presented as black solid lines in Figure 9, top to bottom, respectively. The uncertainty in the estimation of the regression model coefficients is expressed in the form of 95% confidence intervals, indicated by a gray ribbon area around the black solid line. Statistically significant variables are marked with solid red vertical lines.

Simple and multiple linear regression models without regularization both yielded negative association between recent drinking and the right caudal ACC thickness (Figure 8). However, other significant areas differed between simple and multiple regression models with one of the regions (left medial orbital frontal cortex, OFC) exhibiting the positive association with recent drinking. Incorporating the structural connectivity matrix information resulted in three negatively associated regions - right caudal anterior ACC (R-caudACC), left lateral orbitofrontal cortex (L-latOFC), and left precentral gyrus (L-PreCG) for ridge, riPEER and vrPEER estimators (Figure 9). In addition, the cortical thickness of left pars orbitalis showed a trend-level negative association with recent drinking (*p*-value = 0.06 for ridge and riPEER, *p*-value <= 0.07 for vrPEER). By excluding the ridge penalty, the vrPEER method identified a positive association of recent drinking and the right superior frontal gyrus (R-SFG) thickness.

With three regularization models (ridge, riPEER, vrPEER), we found a negative association between cortical thickness of the right caudal ACC and recent drinking, the same region reported by Pennington et al. (2015). A trend-level (*p*-value = 0.06 for Ridge and riPEER; *p*-value = 0.07 for vrPEER) negative association was present in the left caudal ACC thickness. Our finding that the left lateral OFC cortical thickness and recent drinking were negatively related was in agreement with a trend-level cortical thinning of the left OFC in alcohol-only dependent subjects (Pennington et al., 2015). We also observed a negative association of recent drinking and the left precentral gyrus thickness, albeit this area was more posterior than the left middle frontal cortex finding in alcoholics (Nakamura-Palacios et al., 2014), but was homologous to the right precentral gyrus area found to show age effects in alcoholics (Momenan et al., 2012).

None of the brain regions showed positive associations of cortical thickness and recent drinking across all regularization models. The female binge drinkers in Squeglia et al. (2012) exhibited an increased cortical thickness in several frontal areas. The positive association of recent drinking and the SFG cortical thickness in the vrPEER model could reflect a similar relationship in our slightly older male sample. The left parahippocampal gyrus finding in ridge and riPEER models was in unexpected direction although the gray matter volume of that area is known to be affected by the smoking status.

For simplicity, the application of our models was restricted to cortical thickness. It is possible that a more comprehensive picture of the brain morphometry deficits would emerge by also examining other measures, such as cortical surface and gray matter volume. Our regression approach is optimized to model predictors across a wide range of risk for alcoholism rather than focusing on well-defined groups, making direct comparisons more challenging. Finally, we excluded the insula because its structural connectivity information was not available in this analysis.

Importantly, the riPEER estimate mirrors that obtained from ridge. Due to the adaptive properties of riPEER, this result suggests that there is limited information relevant to alcohol consumption in the modularity connectivity graph and its influence in the estimation process is not substantial. From a different perspective, in the riPEER-c with a predefined ordinary ridge regularization parameter, the estimation depends predominantly on the structure imposed by the modularity connectivity graph. This structure can be seen in Figures 9 and 10.

**Figure 9:**
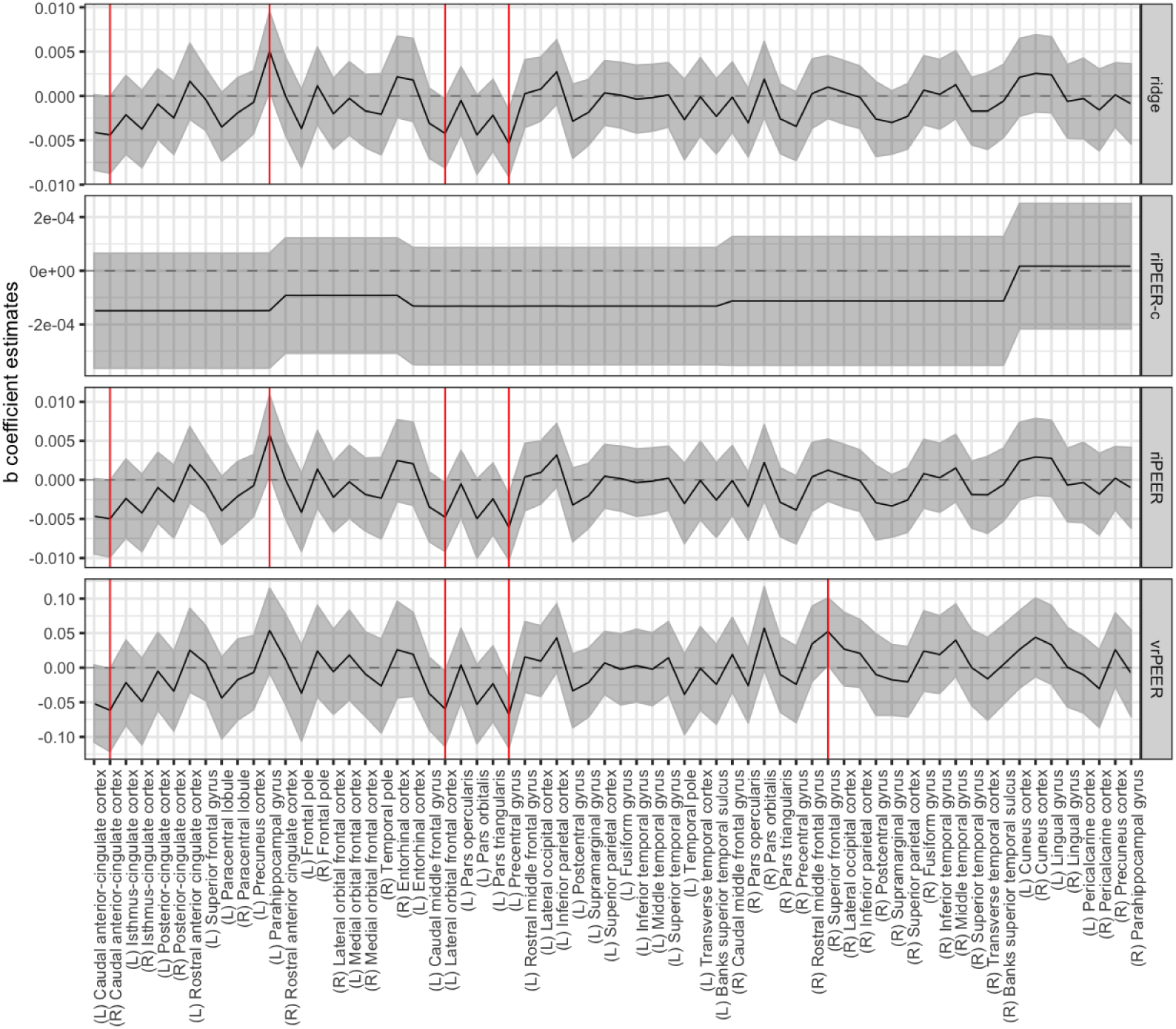
*b̂* coefficient estimates obtained from ridge, riPEER-c, riPEER and vrPEER linear regression estimation methods. Estimates’ values are denoted with a black solid line. 95% confidence intervals are marked with a gray ribbon area. Statistically significant variables are denoted with red vertical solid lines.

**Figure 10:**
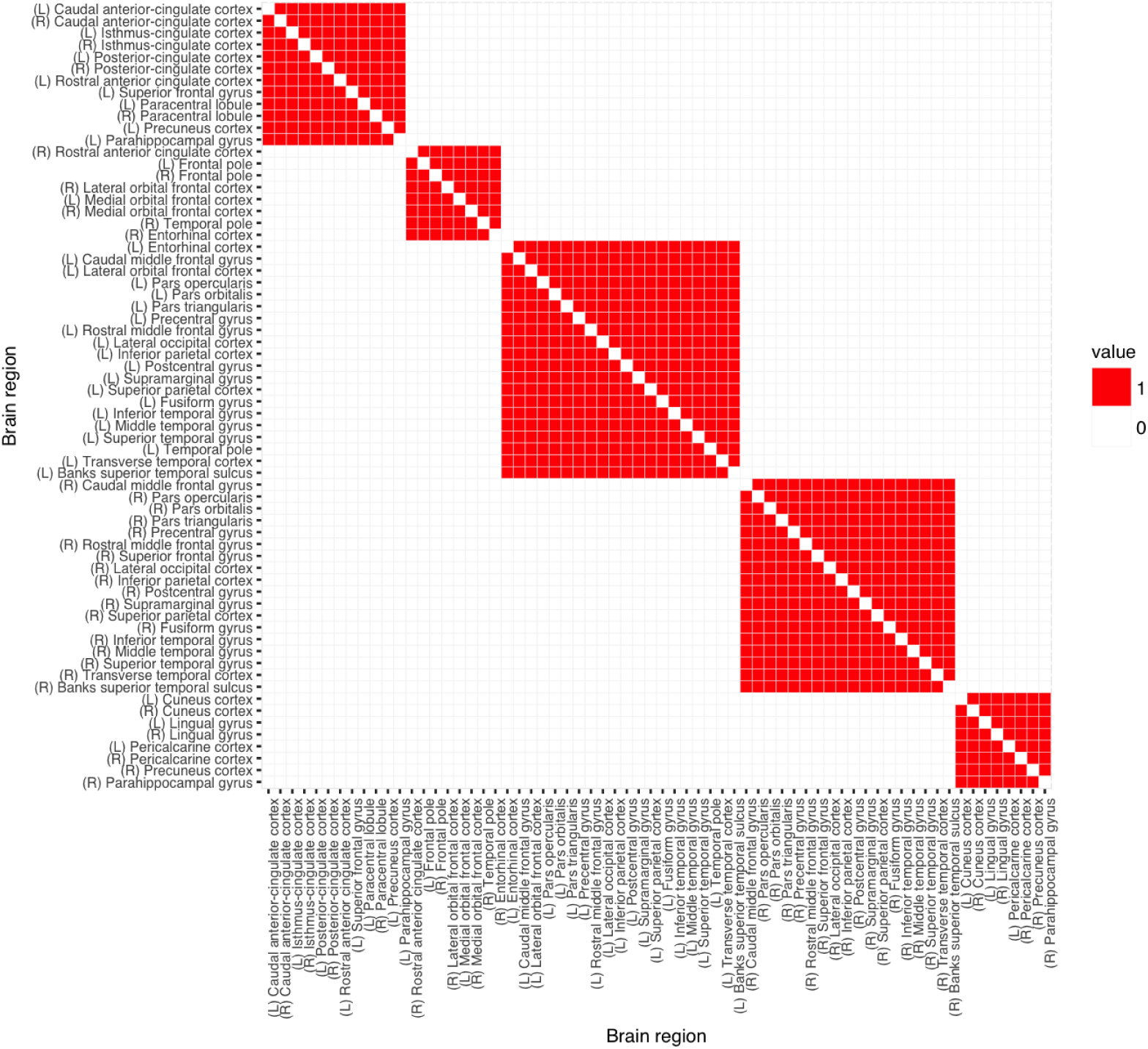
Adjacency matrix of *modularity* graph connectivity information, derived from the connectivity of the brain cortical regions (Sporns, 2013; Cole et al., 2014; Sporns and Betzel, 2016). Each variable belongs to one of five connectivity *modules.*

## 6 Discussion

We have provided a statistically tractable and rigorous way of incorporating external information into the regularized linear model estimation. Our starting point was a generalized ridge or Tikhonov regularization termed PEER Randolph et al. (2012). Here, we concentrate on multi-modal brain imaging data where we obtain structural cortical information from an sMRI scan and structural connectivity information from a DTI scan. We have focused on incorporating the structural connectivity information via the Laplacian-matrix-informed penalty in the Tikhonov regularization framework.

A graph Laplacian matrix is singular and the mixed model equivalency framework requires the invertibility of the penalty matrix. To account for this when using a graph Laplacian-derived penalty term in the PEER framework, we introduced three approaches. Specifically, vrPEER reduces the number of penalized variables by excluding those in the null space of *Q*, while riPEER-c adds a small user-defined multiple of the identity matrix to *Q*. riPEER similarly adds a multiple of the identity to *Q*, but the tuning parameter (i.e., the weight given to the this ridge term) is chosen automatically. Consequently, only riPEER is fully adaptive to the amount of information in *Q* that is relevant to the association between the predictors *Z* and outcome *y*.

Although vrPEER and riPEER-c are more computationally efficient, extensive simulations show that riPEER not only performs better when *Q* contains relevant information, it also performs no worse than ordinary ridge regression when *Q* is not informative.

Application of the proposed methods to study the associations between the alcohol abuse phenotypes and highly correlated cortical thickness predictors using the structural connectivity information resulted in new clinical findings. Specifically, using structural connectivity among 66 cortical regions to define *Q* the proposed methodology found predictive structural imaging markers for the number of drinks per drinking day measure. Negative associations between cortical thickness and Drinks per drinking day were most evident in left and right caudal ACC, left lateral OFC and left precentral gyrus.

Our future work will incorporate additional brain connectivity information arising from other measures of structural brain connectivity including the fractional anisotropy values and fiber length as well as functional connectivity information. We will also combine more than one type of predictor, i.e. use multiple anatomical measures of cortical regions simultaneously.

## Acknowledgements

Research support was partially provided by the NIH grants MH108467, AA017661, AA007611 and AA022476.

## A Appendix

**Figure 11:**
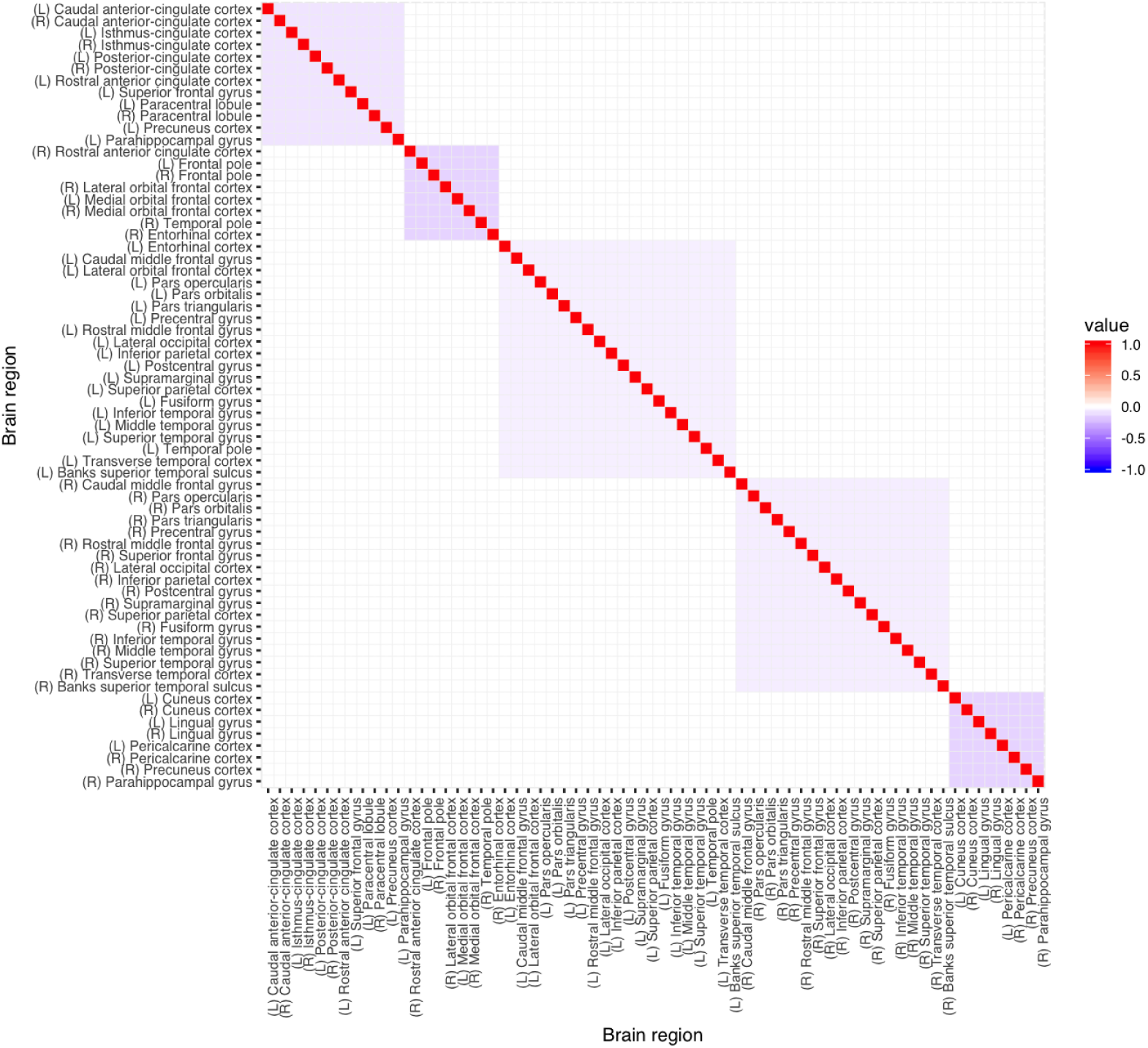
Laplacian matrix of *modularity* graph connectivity information, derived from the connectivity of the brain cortical regions (Sporns, 2013; Cole et al., 2014; Sporns and Betzel, 2016). Each variable belongs to one of five connectivity *modules.*

**Figure 12:**
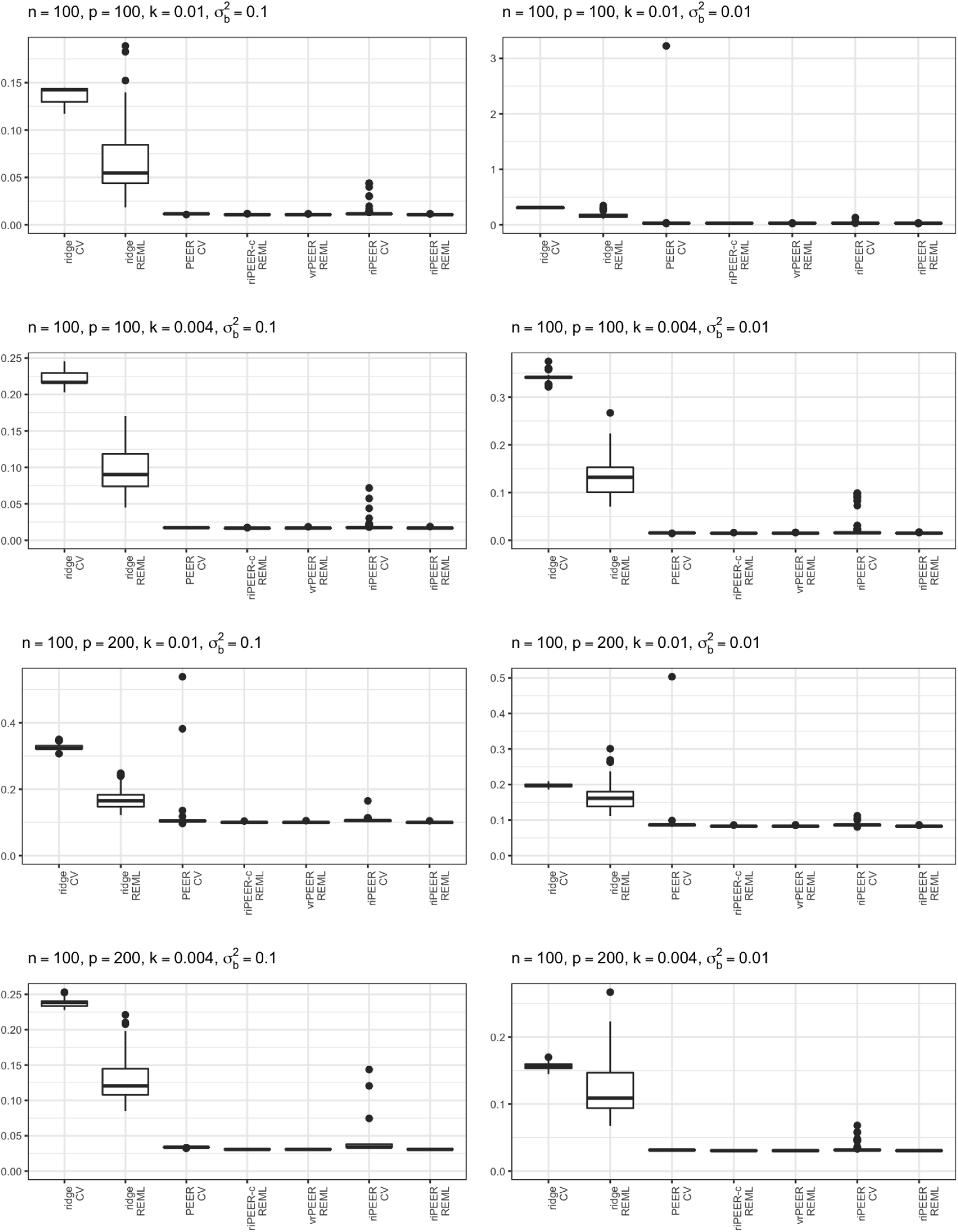
Boxplots of *b* estimation relative error 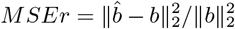 values for different combinations of simulation setup parameters: *n,p,k*,σ^2^_*b*_, in case of informative connectivity information input.

**Figure 13:**
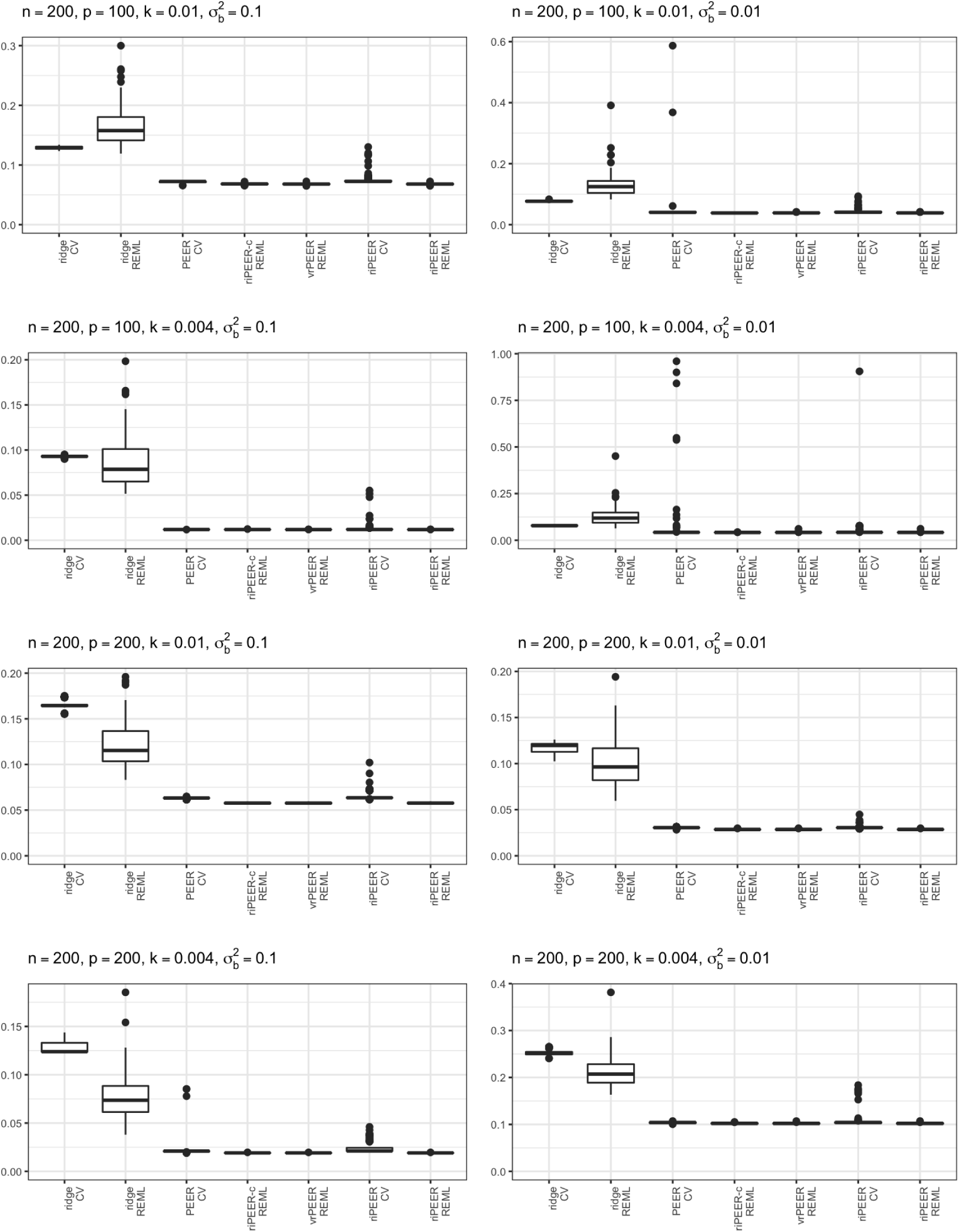
Boxplots of *b* estimation relative error 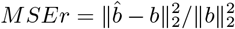 values for different combinations of simulation setup parameters: *n,p,k*,σ^2^_*b*_, in case of informative connectivity information input.

**Table 3:** *b* estimation relative error 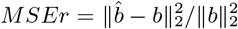 averaged out of 100 experiment runs for each combination of simulation parameters: *n,p,k*,σ^2^_*b*_, steps, in case of partially informative connectivity matrix input, for a graph rewiring performed with the original graphs degree distribution preserved. Minimum estimation *MSEr* values are highlighted (bolded) for each row.

| $n$ | $p$ | $k$ | $\sigma_b^2$ | rewire par.<br>*steps* | ridge<br>REML | riPEER-c<br>REML | vrPEER<br>REML | riPEER<br>REML |
| --- | --- | --- | --- | --- | --- | --- | --- | --- |
| 100 | 100 | 0.004 | 0.01 | 0 | 0.136 | **0.012** | **0.012** | **0.012** |
| 100 | 100 | 0.004 | 0.01 | 10 | 0.127 | **0.014** | 0.015 | 0.015 |
| 100 | 100 | 0.004 | 0.01 | 100 | 0.122 | **0.035** | **0.035** | 0.036 |
| 100 | 100 | 0.004 | 0.01 | 1000 | 0.127 | 0.128 | 0.132 | **0.121** |
| 100 | 100 | 0.004 | 0.01 | 10000 | **0.124** | 0.133 | 0.137 | 0.126 |
| 100 | 100 | 0.004 | 0.1 | 0 | 0.120 | **0.022** | 0.023 | 0.023 |
| 100 | 100 | 0.004 | 0.1 | 10 | 0.120 | **0.025** | **0.025** | **0.025** |
| 100 | 100 | 0.004 | 0.1 | 100 | 0.123 | **0.037** | **0.037** | 0.038 |
| 100 | 100 | 0.004 | 0.1 | 1000 | 0.115 | 0.109 | 0.111 | **0.106** |
| 100 | 100 | 0.004 | 0.1 | 10000 | **0.125** | 0.140 | 0.143 | **0.125** |
| 100 | 100 | 0.01 | 0.01 | 0 | 0.133 | **0.049** | **0.049** | **0.049** |
| 100 | 100 | 0.01 | 0.01 | 10 | 0.140 | 0.050 | 0.050 | 0.050 |
| 100 | 100 | 0.01 | 0.01 | 100 | 0.129 | **0.056** | 0.057 | 0.057 |
| 100 | 100 | 0.01 | 0.01 | 1000 | 0.138 | **0.112** | 0.114 | 0.113 |
| 100 | 100 | 0.01 | 0.01 | 10000 | 0.138 | **0.127** | 0.130 | **0.127** |
| 100 | 100 | 0.01 | 0.1 | 0 | 0.070 | **0.011** | **0.011** | **0.011** |
| 100 | 100 | 0.01 | 0.1 | 10 | 0.066 | **0.011** | **0.011** | **0.011** |
| 100 | 100 | 0.01 | 0.1 | 100 | 0.076 | **0.016** | **0.016** | **0.016** |
| 100 | 100 | 0.01 | 0.1 | 1000 | 0.065 | **0.031** | **0.031** | **0.031** |
| 100 | 100 | 0.01 | 0.1 | 10000 | 0.072 | **0.039** | **0.039** | **0.039** |
| 100 | 200 | 0.004 | 0.01 | 0 | 0.126 | **0.017** | **0.017** | **0.017** |
| 100 | 200 | 0.004 | 0.01 | 10 | 0.132 | **0.017** | **0.017** | **0.017** |
| 100 | 200 | 0.004 | 0.01 | 100 | 0.138 | **0.021** | **0.021** | **0.021** |
| 100 | 200 | 0.004 | 0.01 | 1000 | 0.124 | **0.061** | **0.061** | 0.062 |
| 100 | 200 | 0.004 | 0.01 | 10000 | **0.131** | 0.142 | 0.144 | 0.133 |
| 100 | 200 | 0.004 | 0.1 | 0 | 0.111 | **0.026** | **0.026** | **0.026** |
| 100 | 200 | 0.004 | 0.1 | 10 | 0.107 | **0.026** | **0.026** | **0.026** |
| 100 | 200 | 0.004 | 0.1 | 100 | 0.100 | **0.029** | **0.029** | **0.029** |
| 100 | 200 | 0.004 | 0.1 | 1000 | 0.102 | 0.060 | 0.060 | 0.060 |
| 100 | 200 | 0.004 | 0.1 | 10000 | **0.103** | 0.110 | 0.112 | 0.106 |
| 100 | 200 | 0.01 | 0.01 | 0 | 0.146 | 0.040 | 0.040 | 0.040 |
| 100 | 200 | 0.01 | 0.01 | 10 | 0.152 | 0.040 | 0.040 | 0.040 |
| 100 | 200 | 0.01 | 0.01 | 100 | 0.149 | **0.043** | **0.043** | **0.043** |
| 100 | 200 | 0.01 | 0.01 | 1000 | 0.147 | **0.074** | 0.075 | 0.075 |
| 100 | 200 | 0.01 | 0.01 | 10000 | **0.147** | 0.156 | 0.158 | 0.153 |
| 100 | 200 | 0.01 | 0.1 | 0 | 0.109 | **0.039** | **0.039** | **0.039** |
| 100 | 200 | 0.01 | 0.1 | 10 | 0.106 | **0.039** | **0.039** | **0.039** |
| 100 | 200 | 0.01 | 0.1 | 100 | 0.102 | **0.041** | **0.041** | **0.041** |
| 100 | 200 | 0.01 | 0.1 | 1000 | 0.107 | 0.070 | 0.071 | 0.071 |
| 100 | 200 | 0.01 | 0.1 | 10000 | **0.107** | 0.117 | 0.118 | 0.108 |
| 200 | 100 | 0.004 | 0.01 | 0 | 0.120 | **0.006** | **0.006** | **0.006** |
| 200 | 100 | 0.004 | 0.01 | 10 | 0.125 | **0.007** | **0.007** | **0.007** |
| 200 | 100 | 0.004 | 0.01 | 100 | 0.119 | **0.018** | **0.018** | 0.019 |
| 200 | 100 | 0.004 | 0.01 | 1000 | 0.122 | **0.093** | 0.095 | 0.095 |
| 200 | 100 | 0.004 | 0.01 | 10000 | 0.125 | 0.126 | 0.129 | **0.121** |
| 200 | 100 | 0.004 | 0.1 | 0 | 0.139 | **0.019** | **0.019** | **0.019** |
| 200 | 100 | 0.004 | 0.1 | 10 | 0.140 | **0.021** | **0.021** | **0.021** |
| 200 | 100 | 0.004 | 0.1 | 100 | 0.138 | **0.034** | 0.035 | 0.035 |
| 200 | 100 | 0.004 | 0.1 | 1000 | 0.144 | 0.131 | 0.133 | 0.130 |
| 200 | 100 | 0.004 | 0.1 | 10000 | 0.142 | 0.141 | 0.143 | **0.137** |
| 200 | 100 | 0.01 | 0.01 | 0 | 0.146 | **0.031** | **0.031** | **0.031** |
| 200 | 100 | 0.01 | 0.01 | 10 | 0.142 | **0.033** | **0.033** | **0.033** |
| 200 | 100 | 0.01 | 0.01 | 100 | 0.149 | **0.065** | **0.065** | 0.066 |
| 200 | 100 | 0.01 | 0.01 | 1000 | 0.142 | 0.132 | 0.134 | **0.128** |
| 200 | 100 | 0.01 | 0.01 | 10000 | **0.143** | 0.169 | 0.172 | 0.144 |
| 200 | 100 | 0.01 | 0.1 | 0 | 0.125 | **0.023** | **0.023** | **0.023** |
| 200 | 100 | 0.01 | 0.1 | 10 | 0.123 | **0.027** | **0.027** | **0.027** |
| 200 | 100 | 0.01 | 0.1 | 100 | 0.124 | **0.041** | **0.041** | **0.041** |
| 200 | 100 | 0.01 | 0.1 | 1000 | 0.123 | 0.123 | 0.125 | **0.118** |
| 200 | 100 | 0.01 | 0.1 | 10000 | **0.124** | 0.144 | 0.146 | **0.124** |
| 200 | 200 | 0.004 | 0.01 | 0 | 0.135 | **0.056** | **0.056** | **0.056** |
| 200 | 200 | 0.004 | 0.01 | 10 | 0.135 | **0.055** | **0.055** | **0.055** |
| 200 | 200 | 0.004 | 0.01 | 100 | 0.145 | **0.064** | **0.064** | **0.064** |
| 200 | 200 | 0.004 | 0.01 | 1000 | 0.133 | **0.084** | 0.085 | 0.085 |
| 200 | 200 | 0.004 | 0.01 | 10000 | **0.137** | 0.142 | 0.144 | **0.137** |
| 200 | 200 | 0.004 | 0.1 | 0 | 0.078 | **0.026** | **0.026** | **0.026** |
| 200 | 200 | 0.004 | 0.1 | 10 | 0.076 | **0.026** | **0.026** | **0.026** |
| 200 | 200 | 0.004 | 0.1 | 100 | 0.076 | **0.027** | **0.027** | **0.027** |
| 200 | 200 | 0.004 | 0.1 | 1000 | 0.073 | **0.031** | 0.032 | 0.032 |
| 200 | 200 | 0.004 | 0.1 | 10000 | 0.075 | **0.039** | **0.039** | **0.039** |
| 200 | 200 | 0.01 | 0.01 | 0 | 0.132 | 0.050 | 0.050 | 0.050 |
| 200 | 200 | 0.01 | 0.01 | 10 | 0.124 | 0.050 | 0.050 | 0.050 |
| 200 | 200 | 0.01 | 0.01 | 100 | 0.130 | **0.055** | **0.055** | **0.055** |
| 200 | 200 | 0.01 | 0.01 | 1000 | 0.130 | **0.082** | **0.082** | **0.082** |
| 200 | 200 | 0.01 | 0.01 | 10000 | **0.122** | 0.126 | 0.127 | **0.122** |
| 200 | 200 | 0.01 | 0.1 | 0 | 0.064 | **0.011** | **0.011** | **0.011** |
| 200 | 200 | 0.01 | 0.1 | 10 | 0.065 | **0.011** | **0.011** | **0.011** |
| 200 | 200 | 0.01 | 0.1 | 100 | 0.063 | **0.015** | **0.015** | **0.015** |
| 200 | 200 | 0.01 | 0.1 | 1000 | 0.066 | **0.039** | **0.039** | **0.039** |
| 200 | 200 | 0.01 | 0.1 | 10000 | **0.068** | 0.075 | 0.076 | **0.068** |

**Figure 14:**
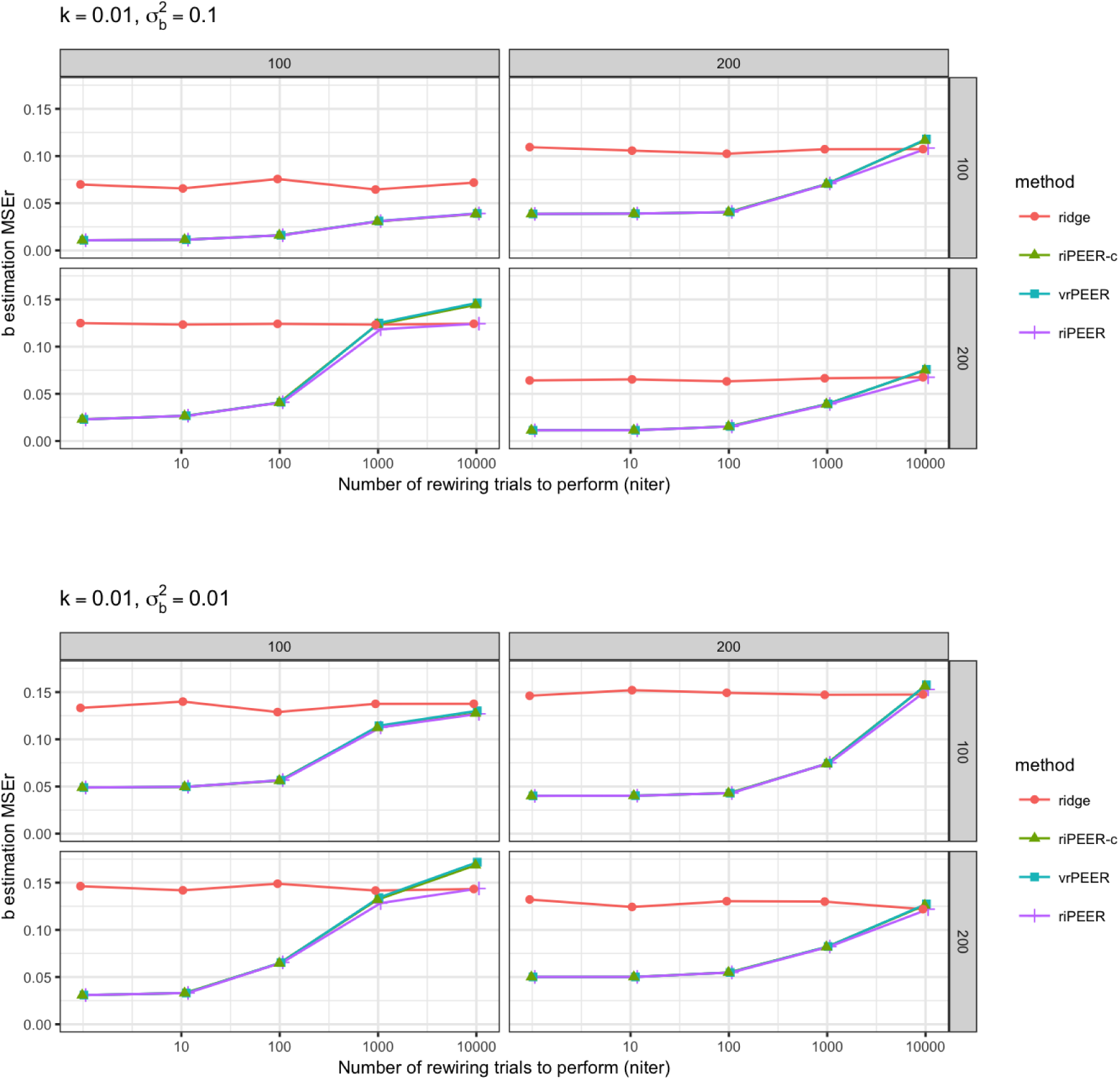
*b* estimation relative error 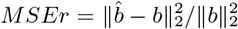 averaged out of 100 experiment runs for different combination of simulation parameters: *k*σ^2^_*b*_, *steps* ∈ {0,10,100,1000,1000} (denoted by x-axis labels), for *p* = 100 and 200 (left-side and right-side panels, respectively), *n* = 100 and 200 (upper-side and bottom-side panels, respectively), in case of partially informative connectivity matrix input. A graph rewiring performed with the original graphs degree distribution preserved was used. Minimum estimation *MSEr* values are highlighted (bolded) for each row.

**Figure 15:**
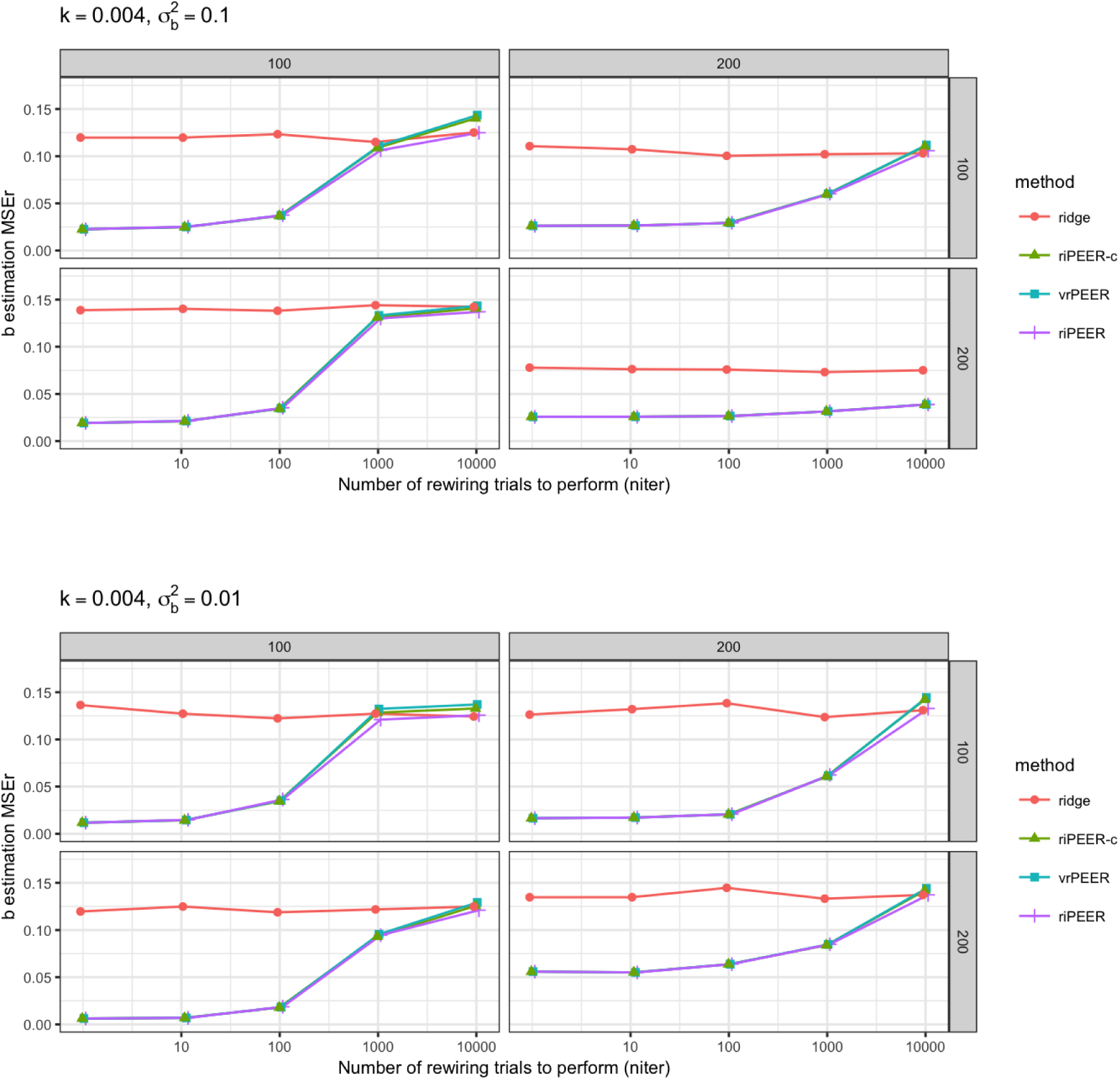
*b* estimation relative error 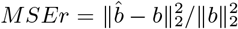 averaged out of 100 experiment runs for different combination of simulation parameters: *k*σ^2^_*b*_, *steps* ∈ {0,10,100,1000,1000} (denoted by x-axis labels), for *p* = 100 and 200 (left-side and right-side panels, respectively), *n* = 100 and 200 (upper-side and bottom-side panels, respectively), in case of partially informative connectivity matrix input. A graph rewiring performed with the original graphs degree distribution preserved was used. Minimum estimation *MSEr* values are highlighted (bolded) for each row.

**Table 4:** *b* estimation relative error 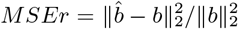 averaged out of 100 experiment runs for each combination of simulation parameters: *n,p,k*,σ^2^_*b*_, *prob*, in case of partially informative connectivity matrix input, for a graph rewiring performed with a constant probability for a graph node to be rewired. Minimum estimation *MSEr* values are highlighted (bolded) for each row.

| $n$ | $p$ | $k$ | $\sigma_b^2$ | rewire par.<br>$prob$ | ridge<br>REML | riPEER-c<br>REML | vrPEER<br>REML | riPEER<br>REML |
| --- | --- | --- | --- | --- | --- | --- | --- | --- |
| 100 | 100 | 0.004 | 0.01 | 0 | 0.167 | **0.019** | **0.019** | **0.019** |
| 100 | 100 | 0.004 | 0.01 | 0.2 | 0.170 | **0.087** | 0.089 | 0.090 |
| 100 | 100 | 0.004 | 0.01 | 0.4 | 0.170 | **0.146** | 0.151 | 0.151 |
| 100 | 100 | 0.004 | 0.01 | 0.6 | 0.169 | **0.163** | 0.167 | 0.169 |
| 100 | 100 | 0.004 | 0.01 | 0.8 | **0.165** | 0.205 | 0.210 | **0.165** |
| 100 | 100 | 0.004 | 0.01 | 1 | **0.172** | 0.196 | 0.201 | 0.173 |
| 100 | 100 | 0.004 | 0.1 | 0 | 0.095 | **0.021** | **0.021** | **0.021** |
| 100 | 100 | 0.004 | 0.1 | 0.2 | 0.091 | **0.053** | 0.054 | 0.054 |
| 100 | 100 | 0.004 | 0.1 | 0.4 | 0.098 | **0.082** | 0.084 | **0.082** |
| 100 | 100 | 0.004 | 0.1 | 0.6 | 0.095 | **0.083** | 0.086 | **0.083** |
| 100 | 100 | 0.004 | 0.1 | 0.8 | **0.091** | 0.104 | 0.106 | 0.101 |
| 100 | 100 | 0.004 | 0.1 | 1 | **0.096** | 0.105 | 0.108 | 0.102 |
| 100 | 100 | 0.01 | 0.01 | 0 | 0.120 | **0.006** | **0.006** | **0.006** |
| 100 | 100 | 0.01 | 0.01 | 0.2 | 0.120 | **0.057** | 0.058 | 0.060 |
| 100 | 100 | 0.01 | 0.01 | 0.4 | 0.118 | **0.101** | 0.103 | **0.101** |
| 100 | 100 | 0.01 | 0.01 | 0.6 | 0.126 | 0.134 | 0.137 | **0.122** |
| 100 | 100 | 0.01 | 0.01 | 0.8 | **0.123** | 0.158 | 0.162 | 0.124 |
| 100 | 100 | 0.01 | 0.01 | 1 | **0.121** | 0.147 | 0.150 | 0.123 |
| 100 | 100 | 0.01 | 0.1 | 0 | 0.070 | **0.011** | **0.011** | **0.011** |
| 100 | 100 | 0.01 | 0.1 | 0.2 | 0.063 | **0.028** | **0.028** | **0.028** |
| 100 | 100 | 0.01 | 0.1 | 0.4 | 0.067 | **0.035** | **0.035** | **0.035** |
| 100 | 100 | 0.01 | 0.1 | 0.6 | 0.071 | **0.031** | **0.031** | **0.031** |
| 100 | 100 | 0.01 | 0.1 | 0.8 | 0.061 | **0.037** | 0.038 | 0.038 |
| 100 | 100 | 0.01 | 0.1 | 1 | 0.072 | **0.038** | **0.038** | **0.038** |
| 100 | 200 | 0.004 | 0.01 | 0 | 0.117 | **0.023** | **0.023** | **0.023** |
| 100 | 200 | 0.004 | 0.01 | 0.2 | 0.115 | **0.061** | 0.062 | 0.062 |
| 100 | 200 | 0.004 | 0.01 | 0.4 | 0.110 | **0.096** | 0.098 | 0.097 |
| 100 | 200 | 0.004 | 0.01 | 0.6 | 0.122 | **0.106** | 0.108 | 0.107 |
| 100 | 200 | 0.004 | 0.01 | 0.8 | **0.120** | 0.122 | 0.124 | **0.120** |
| 100 | 200 | 0.004 | 0.01 | 1 | **0.117** | 0.122 | 0.124 | 0.120 |
| 100 | 200 | 0.004 | 0.1 | 0 | 0.086 | **0.018** | **0.018** | **0.018** |
| 100 | 200 | 0.004 | 0.1 | 0.2 | 0.090 | **0.048** | 0.049 | 0.049 |
| 100 | 200 | 0.004 | 0.1 | 0.4 | 0.090 | **0.079** | 0.080 | **0.079** |
| 100 | 200 | 0.004 | 0.1 | 0.6 | 0.091 | 0.091 | 0.092 | **0.089** |
| 100 | 200 | 0.004 | 0.1 | 0.8 | **0.090** | 0.092 | 0.093 | **0.090** |
| 100 | 200 | 0.004 | 0.1 | 1 | **0.089** | 0.093 | 0.094 | 0.091 |
| 100 | 200 | 0.01 | 0.01 | 0 | 0.139 | **0.037** | **0.037** | **0.037** |
| 100 | 200 | 0.01 | 0.01 | 0.2 | 0.140 | **0.090** | 0.091 | 0.091 |
| 100 | 200 | 0.01 | 0.01 | 0.4 | 0.139 | **0.119** | 0.121 | 0.121 |
| 100 | 200 | 0.01 | 0.01 | 0.6 | **0.144** | 0.147 | 0.149 | 0.145 |
| 100 | 200 | 0.01 | 0.01 | 0.8 | **0.138** | 0.142 | 0.144 | **0.138** |
| 100 | 200 | 0.01 | 0.01 | 1 | **0.137** | 0.145 | 0.147 | 0.139 |
| 100 | 200 | 0.01 | 0.1 | 0 | 0.098 | **0.033** | **0.033** | **0.033** |
| 100 | 200 | 0.01 | 0.1 | 0.2 | 0.103 | **0.066** | **0.066** | 0.067 |
| 100 | 200 | 0.01 | 0.1 | 0.4 | 0.106 | **0.088** | 0.089 | 0.089 |
| 100 | 200 | 0.01 | 0.1 | 0.6 | 0.104 | **0.100** | 0.102 | 0.101 |
| 100 | 200 | 0.01 | 0.1 | 0.8 | 0.104 | **0.103** | 0.104 | **0.103** |
| 100 | 200 | 0.01 | 0.1 | 1 | **0.106** | 0.113 | 0.115 | 0.112 |
| 200 | 100 | 0.004 | 0.01 | 0 | 0.087 | **0.017** | **0.017** | **0.017** |
| 200 | 100 | 0.004 | 0.01 | 0.2 | 0.099 | **0.045** | 0.046 | 0.046 |
| 200 | 100 | 0.004 | 0.01 | 0.4 | 0.093 | **0.060** | 0.061 | **0.060** |
| 200 | 100 | 0.004 | 0.01 | 0.6 | 0.094 | **0.066** | 0.067 | **0.066** |
| 200 | 100 | 0.004 | 0.01 | 0.8 | 0.094 | **0.062** | 0.063 | 0.063 |
| 200 | 100 | 0.004 | 0.01 | 1 | 0.100 | **0.056** | 0.058 | 0.057 |
| 200 | 100 | 0.004 | 0.1 | 0 | 0.080 | **0.009** | **0.009** | **0.009** |
| 200 | 100 | 0.004 | 0.1 | 0.2 | 0.083 | **0.043** | **0.043** | **0.043** |
| 200 | 100 | 0.004 | 0.1 | 0.4 | 0.085 | **0.059** | 0.061 | 0.060 |
| 200 | 100 | 0.004 | 0.1 | 0.6 | 0.078 | 0.073 | 0.074 | **0.072** |
| 200 | 100 | 0.004 | 0.1 | 0.8 | **0.081** | 0.088 | 0.089 | 0.086 |
| 200 | 100 | 0.004 | 0.1 | 1 | **0.084** | 0.091 | 0.093 | 0.089 |
| 200 | 100 | 0.01 | 0.01 | 0 | 0.183 | **0.038** | **0.038** | **0.038** |
| 200 | 100 | 0.01 | 0.01 | 0.2 | 0.180 | **0.100** | 0.101 | 0.102 |
| 200 | 100 | 0.01 | 0.01 | 0.4 | 0.181 | 0.179 | 0.182 | **0.175** |
| 200 | 100 | 0.01 | 0.01 | 0.6 | 0.179 | 0.191 | 0.195 | **0.178** |
| 200 | 100 | 0.01 | 0.01 | 0.8 | **0.177** | 0.195 | 0.198 | 0.178 |
| 200 | 100 | 0.01 | 0.01 | 1 | **0.180** | 0.204 | 0.207 | 0.181 |
| 200 | 100 | 0.01 | 0.1 | 0 | 0.115 | **0.033** | **0.033** | **0.033** |
| 200 | 100 | 0.01 | 0.1 | 0.2 | 0.126 | **0.092** | 0.094 | 0.094 |
| 200 | 100 | 0.01 | 0.1 | 0.4 | 0.118 | **0.100** | 0.102 | **0.100** |
| 200 | 100 | 0.01 | 0.1 | 0.6 | **0.116** | 0.141 | 0.143 | 0.127 |
| 200 | 100 | 0.01 | 0.1 | 0.8 | **0.117** | 0.153 | 0.155 | 0.136 |
| 200 | 100 | 0.01 | 0.1 | 1 | **0.128** | 0.140 | 0.143 | 0.129 |
| 200 | 200 | 0.004 | 0.01 | 0 | 0.162 | **0.070** | **0.070** | **0.070** |
| 200 | 200 | 0.004 | 0.01 | 0.2 | 0.172 | **0.122** | 0.123 | 0.123 |
| 200 | 200 | 0.004 | 0.01 | 0.4 | 0.162 | **0.144** | 0.145 | **0.144** |
| 200 | 200 | 0.004 | 0.01 | 0.6 | 0.158 | 0.158 | 0.160 | **0.157** |
| 200 | 200 | 0.004 | 0.01 | 0.8 | **0.163** | 0.164 | 0.166 | **0.163** |
| 200 | 200 | 0.004 | 0.01 | 1 | **0.169** | 0.170 | 0.172 | **0.169** |
| 200 | 200 | 0.004 | 0.1 | 0 | 0.081 | **0.032** | **0.032** | **0.032** |
| 200 | 200 | 0.004 | 0.1 | 0.2 | 0.086 | **0.038** | **0.038** | **0.038** |
| 200 | 200 | 0.004 | 0.1 | 0.4 | 0.081 | **0.040** | **0.040** | **0.040** |
| 200 | 200 | 0.004 | 0.1 | 0.6 | 0.084 | **0.041** | **0.041** | **0.041** |
| 200 | 200 | 0.004 | 0.1 | 0.8 | 0.079 | **0.041** | **0.041** | **0.041** |
| 200 | 200 | 0.004 | 0.1 | 1 | 0.081 | **0.042** | 0.043 | 0.043 |
| 200 | 200 | 0.01 | 0.01 | 0 | 0.113 | **0.040** | **0.040** | **0.040** |
| 200 | 200 | 0.01 | 0.01 | 0.2 | 0.114 | **0.067** | 0.068 | 0.068 |
| 200 | 200 | 0.01 | 0.01 | 0.4 | 0.112 | **0.079** | **0.079** | 0.080 |
| 200 | 200 | 0.01 | 0.01 | 0.6 | 0.116 | **0.093** | 0.094 | 0.094 |
| 200 | 200 | 0.01 | 0.01 | 0.8 | 0.115 | **0.099** | 0.100 | **0.099** |
| 200 | 200 | 0.01 | 0.01 | 1 | 0.116 | **0.092** | 0.093 | 0.093 |
| 200 | 200 | 0.01 | 0.1 | 0 | 0.142 | **0.043** | **0.043** | **0.043** |
| 200 | 200 | 0.01 | 0.1 | 0.2 | 0.142 | **0.092** | **0.092** | 0.093 |
| 200 | 200 | 0.01 | 0.1 | 0.4 | 0.138 | **0.120** | 0.121 | **0.120** |
| 200 | 200 | 0.01 | 0.1 | 0.6 | **0.141** | 0.142 | 0.143 | **0.141** |
| 200 | 200 | 0.01 | 0.1 | 0.8 | **0.142** | 0.144 | 0.145 | 0.143 |
| 200 | 200 | 0.01 | 0.1 | 1 | **0.140** | 0.150 | 0.151 | 0.147 |

**Figure 16:**
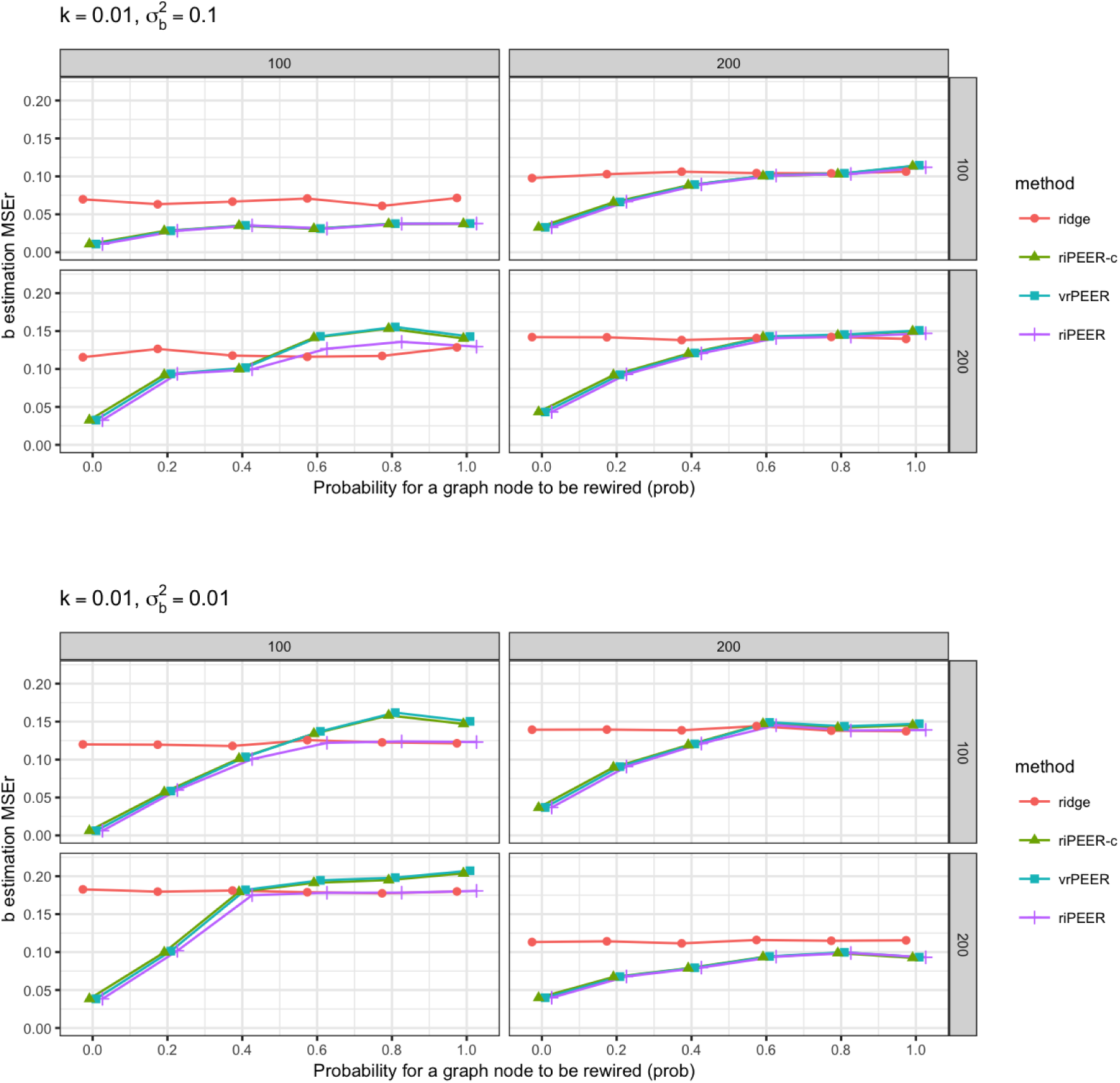
*b* estimation relative error 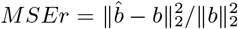 averaged out of 100 experiment runs for different combination of simulation parameters: *k*,σ^2^_*b*_, *prob* ∈ {0, 0.2, 0.4, 0.6, 0.8,1} (denoted by x-axis labels), for *p* = 100 and 200 (left-side and right-side panels, respectively), *n* = 100 and 200 (upper-side and bottom-side panels, respectively), in case of partially informative connectivity matrix input. A graph rewiring with a constant probability for a graph node to be rewired was used. Minimum estimation *MSEr* values are highlighted (bolded) for each row.

**Figure 17:**
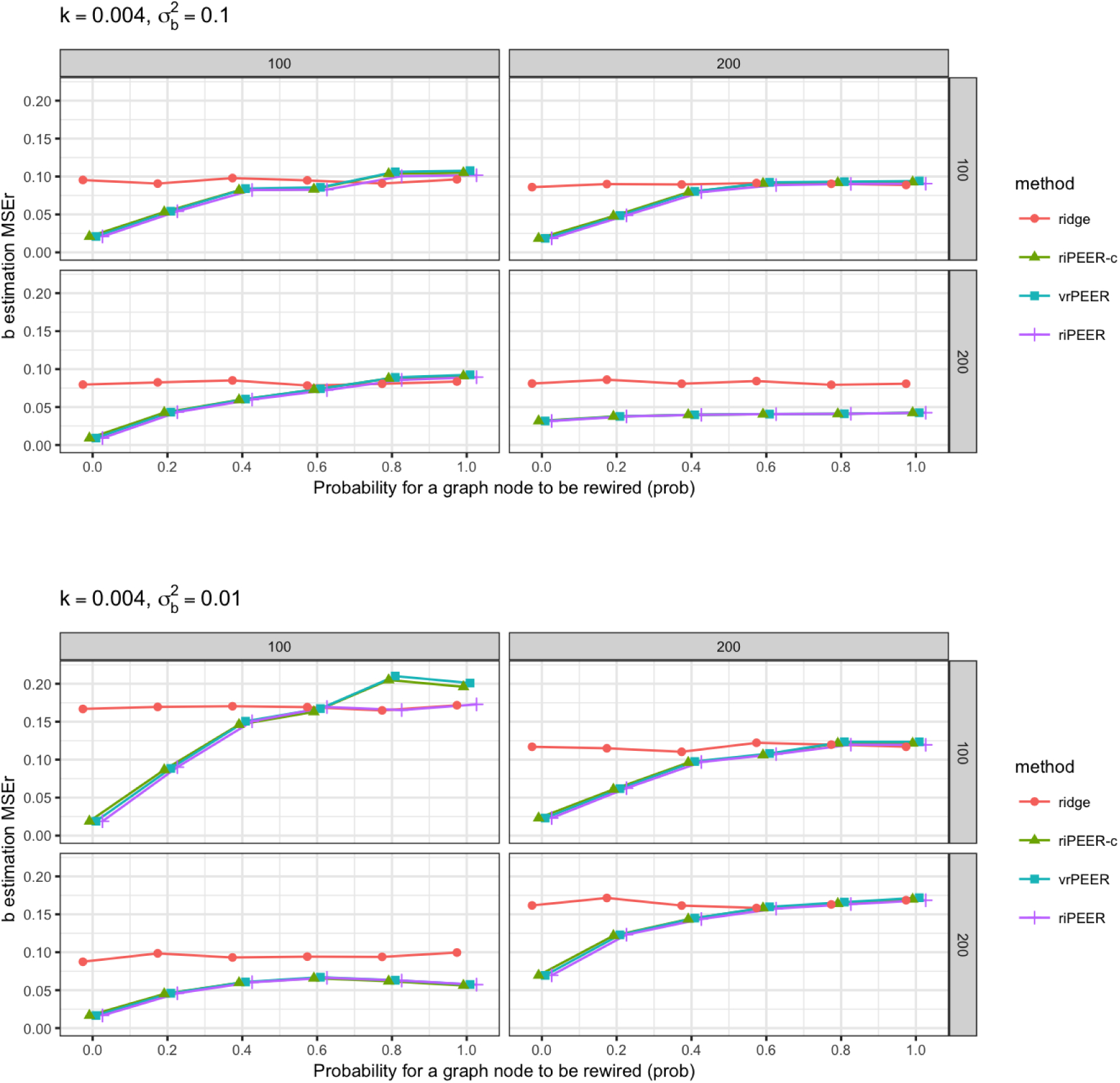
*b* estimation relative error 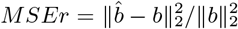 averaged out of 100 experiment runs for different combination of simulation parameters: *k*,σ^2^_*b*_, *prob* ∈ {0, 0.2, 0.4, 0.6, 0.8,1} (denoted by x-axis labels), for *p* = 100 and 200 (left-side and right-side panels, respectively), *n* = 100 and 200 (upper-side and bottom-side panels, respectively), in case of partially informative connectivity matrix input. A graph rewiring with a constant probability for a graph node to be rewired was used. Minimum estimation *MSEr* values are highlighted (bolded) for each row.

**Figure 18:**
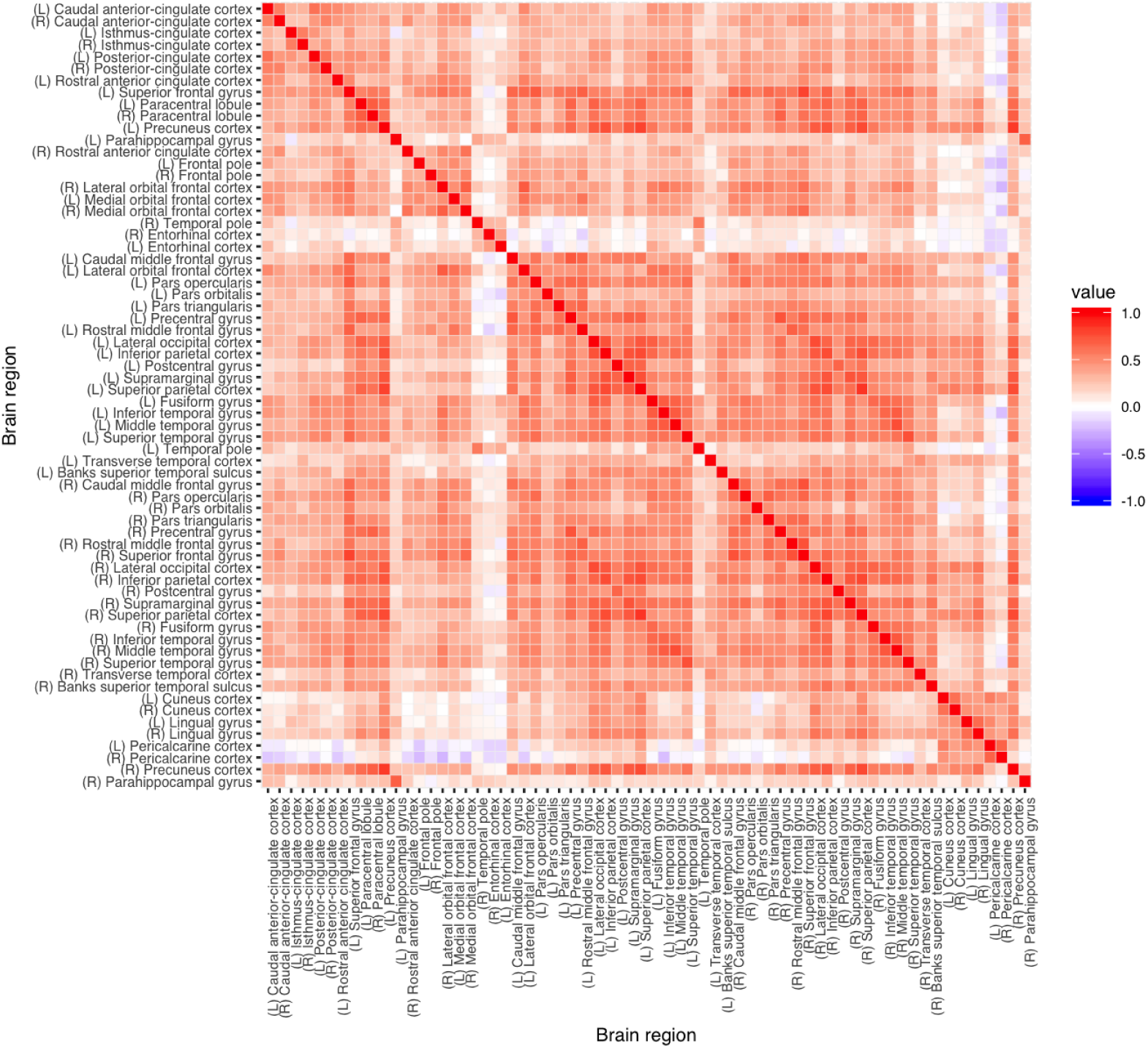
Pairwise Pearson correlation coefficient for cortical average thickness measurements of 66 brain regions of 148 study subjects.

